# Behavioral characterization of dynamic facial expression perception in rhesus monkeys *(Macaca mulatta)* using naturalistic and synthetic stimuli

**DOI:** 10.64898/2026.03.18.712749

**Authors:** Ramona Siebert, Nick Taubert, Martin A. Giese, Peter Thier

**Affiliations:** Cognitive Neurology Laboratory, Hertie Institute for Clinical Brain Research, University of Tübingen, Germany; Computational Sensomotorics, Centre for Integrative Neuroscience & Hertie Institute for Clinical Brain Research, University of Tübingen, Germany; Graduate School of Neural and Behavioural Sciences, University of Tübingen, Germany

**Author notes:** equal contributions (Martin A. Giese, Peter Thier).

**Keywords:** Facial expression, rhesus macaque, social cognition, face avatar

## Abstract

Facial displays are an important channel of communication for primates, yet it remains unclear based on which criteria monkeys evaluate facial expressions. We trained two rhesus macaques to categorize videos of four facial expression types (neutral, lip-smacking, silent bared-teeth (SBT) and open-mouth threat displays) and then tested generalization to novel individuals and avatars while measuring pupillary responses. The animals were able to categorize facial expressions and generalize their classifications to novel, untrained videos, although performance was not perfect. Misclassifications were not driven solely by visual similarity between expression categories, as demonstrated by a novel automated approach integrating deep learning-based motion tracking with the Macaque Facial Action Coding System to enable objective quantification of facial action units. Instead, open-mouth threats were readily categorized correctly and distinguished from lip-smacking, while also eliciting the strongest arousal responses. By contrast, categorization of SBT displays varied substantially across stimulus identities, influenced by body weight, gaze direction, and coordinated movements of eyebrows and ears. Morphed avatar expressions were categorized according to expression component intensity, demonstrating graded perception. Avatar manipulations revealed that categorization was robust against lack of coherent motion, lack of realism beyond a basic level of texture, and to some extent transcended species form boundaries to a human face. However, human facial expressions elicited no systematic categorizations or differential arousal. Our findings indicate that conspecific facial expressions were perceived as functionally meaningful, context-dependent social signals shaped by all components of facial behavior and signaler characteristics, rather than fixed, mouth movement-defined categories.

## INTRODUCTION

The ability to distinguish and interpret facial expressions is an important social skill for all primate species. Since Darwin (1872) pioneered the systematic investigation of facial displays in humans and animals, many ethological studies in non-human primates have provided descriptions of facial movements and the situations in which they usually occur (e.g. Andrew, 1963; Hinde, 1966; Maestripieri, 1997; Partan, 2002; Preuschoft, 2000; van Hooff, 1967). Laboratory studies have reported how macaque monkeys visually scan facials displays (Gothard, Erickson, & Amaral, 2004; Mosher, Zimmerman, & Gothard, 2011; Nahm, Perret, Amaral, & Albright, 1997; Siebert et al., 2020) and examined neural responses to viewing conspecific facial expressions (e.g. Gothard, Battaglia, Erickson, Spitler, & Amaral, 2007; Hadj-Bouziane, Bell, Knusten, Ungerleider, & Tootell, 2008; Hasselmo, Rolls, & Baylis, 1989; Hoffman, Gothard, Schmid, & Logothetis, 2007; J. Taubert et al., 2020; Yang & Freiwald, 2021; Zhu et al., 2013). The most frequently employed expressions in these studies were aggressive open-mouth threat, affiliative/appeasing lip-smacking with puckered lips and rapid opening and closing of the mouth, and fearful/submissive fear-grin or -grimace, more objectively called the silent bared-teeth display (SBT), with mouth corners and lips retracted to expose the teeth. However, despite the widespread use of these stereotypical categories, few studies have explicitly asked monkeys to behaviorally discriminate facial expressions. Kanazawa (1996) reported that one Japanese macaque differentiated facial displays in a match-to-sample paradigm along a “subordinate – dominant” dimension, distinguishing SBT and threat faces by mouth shape and eye width, and a “neutral – tense” dimension, separating relaxed faces from faces with raised eyebrows, whereas the macaque only distinguished the human smile among other human expressions. This study only used one Japanese macaque’s and one human’s displays as stimuli, leaving unclear whether the subject resorted to any semantic information about expression categories or simply differentiated dissimilar pictures. Parr and Heintz (2009) performed a similar experiment with seven rhesus macaques and the categories open-mouth threat, SBT, play face and scream face. The macaques struggled when the identity of the sample, the matching and the non-matching stimulus were all different and confusion apparently occurred due to shared features between expression categories, especially mouth opening and teeth exposure. However, overlapping features alone could not explain the overall error pattern. Notably, the macaques exhibited a bias for selecting the play face, presumably the most positively valued among the offered categories. An analogous approach was used by Micheletta, Whitehouse, Parr, and Waller (2015) who tested three crested macaques and reported that the visual similarity between expressions was not correlated with performance, and speculated that functional similarity between expressions could be responsible for errors. J. Taubert and Japee (2024) trained four rhesus macaques to select a “positive” facial expression (neutral or lip-smack) over a “negative” expression (threat or SBT). When tested with novel images, the macaques most frequently selected the lip-smack stimuli and least frequently selected the threat stimuli, as expected, but also selected two SBT displays over most neutral faces, while the other three SBTs ranked among the threat displays, suggesting variable valence of SBTs. All four studies relied on static images, although Micheletta et al. (2015) also included dynamic video stimuli and anecdotally noted improved performance.

While these studies have demonstrated that macaques are able to distinguish facial expressions in experimental settings, it remains unclear I) which strategy they used to discriminate expressions and II) which facial motion, shape and appearance features are necessary and sufficient for the perception and differentiation of a facial expression category. To answer these questions, we trained two rhesus macaques to explicitly discriminate videos of neutral, lip-smacking, SBT and threat displays in a forced-choice categorization paradigm, avoiding a potential bias to select more pleasant expressions in a match-to-sample configuration. We subsequently assessed them using novel expression videos in catch trials and measured physiological arousal through pupillary responses. For part I), the test videos were filmed real macaques and a naturalistic monkey avatar. We reasoned that if the macaques used elementary facial features to make their decisions without attaching any meaning to the videos, they would perform comparably well across expression categories for the new test videos, with mistakes occurring among visually similar expressions and without physiological reactions to the videos. However, if the macaques perceived the videos as the described mouth movement-defined morphological categories (neutral, lip-smacking, SBT and open-mouth threat), each with discrete meaning, they would exhibit the same performance and error pattern, but experience differential arousal when seeing the different expression categories. To quantify similarity among the expressions, we combined the Facial Action Coding System (FACS), which allows the objective, muscle-based description of facial movements (Ekman & Friesen, 1978; Ekman, Friesen, & Hager, 2002; Parr, Waller, Burrows, Gothard, & Vick, 2010) and deep learning-based motion tracking (DeepLabCut) (Mathis et al., 2018; Nath et al., 2019). If the meaning the macaques extracted from the stimuli is not fully captured by the defined categories, inconsistent performance across expression categories and test identities should occur and visual similarity should fail to explain the pattern of miscategorizations. Instead confusions between expressions with similar function should occur, like lip-smacking and neutral, lip-smacking and SBT or SBT and threat, and a clear distinction between lip-smacking and threat. However, if they only learned the association of the rewarded target for each individual training video, they should fail to categorize the new videos according to expression type, although rhesus macaques were able to categorize action videos in a similar paradigm (Cui et al., 2023). For part II), we made use of a monkey avatar to continuously morph one expression into another, to remove motion coherence, to systematically change the facial appearance by gradually decreasing realism (no fur, no color, no texture, no shape), to change the species-specific facial form to a human avatar and to change the species-specific motion pattern to human expressions. Note that we use the conventional term facial expression interchangeably with facial display, facial configuration, facial behavior or facial movement pattern, without assuming any expression of an emotional state.

## MATERIALS AND METHODS

### Subjects

Data was collected from two adult male rhesus macaques (*Macaca mulatta*), monkey 1 (7 years, 10.5 kg) and monkey 2 (8 years, 8.5 kg). They were born at the German Primate Center, raised in social groups, and pair-housed in our laboratory with visual and auditory contact to conspecifics. For both monkeys, this was the first scientific experiment in which they participated. Following chair training, they were implanted with individually adapted titanium head posts for head immobilization during eye tracking and subsequently planned neurophysiological experiments. All surgical procedures were conducted aseptically under general anesthesia and control of vital parameters. After recovery, the monkeys were gradually trained to accept the head immobilization and to receive liquid rewards (juice and water) for fixating dots and images on a computer screen. All animal procedures were approved by the veterinary administration (Regierungspräsidium Tübingen, Abteilung Tierschutz) and complied with German law (Tierschutz-Versuchstierverordnung) and the National Institutes of Health’s Guide for the Care and Use of Laboratory Animals.

### Visual stimuli

#### Real monkeys

Training stimuli consisted of 13 videos per expression category from diverse sources: 13 rhesus macaques from our and neighboring laboratories (known and unknown to subjects) sitting in a primate chair with their heads free to move, natural environment footage (www.youtube.com/@rhesusvideo6831, channel photo credit: Lauren Brent), and still photographs (PrimFace database: http://visiome.neuroinf.jp/primface/, funded by Grant-in-Aid for Scientific research on Innovative Areas, “Face Perception and Recognition” from the Ministry of Education, Culture, Sports, Science, and Technology (MEXT), Japan). This provided variability in background, face size, and dynamic content. The test stimuli (all male rhesus macaques) comprised one set of expression videos of monkeys contained in the training set (unknown individuals) which had not been shown during training, and nine new identities. Eight of these (ID 1–2, 5–9) were monkeys from neighboring laboratories (ages 12–17, weight 8.8–15.6 kg), unknown to the subjects, and one corresponded to the subject’s cage partner (ID 3 and 4, respectively). Four test expression sets (ID 1–4) comprised the complete desired expression repertoire (neutral, lip-smacking, silent bared-teeth and open-mouth threat), whereas five individuals (ID 5–9) did not show the silent bared-teeth display. Facial behaviors were elicited by the experimenter interacting with the monkey, mimicking macaque displays and showing positive (treats) and negative (tools) stimuli. Each test monkey was filmed in a single recording session under standardized conditions seated in a primate chair with unrestrained head against a neutral white background. Test videos were cropped to show the heads only, cut into clips of single continuous facial displays, manually scored according to the Macaque Facial Action Coding System (MaqFACS) (Parr et al., 2010) and assigned to four expression categories. “Neutral”: no activation of mouth action units (AUs) and little to no eyebrow or ear movements. “Lip-smacking”: AU18i (true lip pucker) and AD181 (lip-smacking action descriptor). “Silent bared-teeth” (SBT) display (fear-grin): AU10 (upper lip raiser), AU12 (lip corner puller), AU16 (lower lip depressor), AU25 (lips parted) and optionally AU26 (jaw drop). “Open-mouth threat”: AU25 (lips parted) and either AU26 (jaw drop) (without any AU12) or AU27 (mouth stretch). Only clear expressions were used. Similar definitions for macaque facial expression categories were employed by others (Parr et al., 2010; J. Taubert & Japee, 2024).

#### Monkey and human avatars

The monkey avatar was created from an MRI-based mesh surface controlled by ribbons modeled after rhesus macaque facial muscles, driven by motion-captured facial movements (Siebert et al., 2020; N. Taubert et al., 2021). We have previously demonstrated that this monkey avatar elicits behavioral reactions similar to real macaque videos and does not fall into the “uncanny valley” of aversion in the visual perception of rhesus macaques towards synthetic agents (Siebert et al., 2020). The human avatar used equivalent control points following human muscle anatomy. In addition to the pure, prototypical motion-captured expressions, the monkey avatar displayed expressions morphed in 25 % intensity steps between prototypical category pairs. Additional test conditions included variations of the prototypical expressions monkey avatar: temporally scrambled frames, point-light displays (movement patterns without texture) and degraded realism versions (furless, grayscale, wireframe); as well as human expressions (smiling/happy, gasping/fearful, scowling/angry) on human and monkey avatars. More details on the avatar stimuli are available in the Supplementary Methods.

### Experimental paradigm and procedure

Subjects sat head-restrained in a primate chair 60 cm in front of a computer monitor in a darkened booth. Stimuli were presented via the NREC open-source control system (https://nrec.neurologie.uni-tuebingen.de/) on a gray background. Eye position and pupil size (vertical diameter) were recorded with an eye tracker (ISCAN ETL-200, 120 Hz). A trial was initiated by fixation on a dot (radius 0.2° visual angle) for 1 s, followed by a 1–2 s centrally presented video (12° x 12°) and 0.5 s of blank screen. Then four differently colored targets (0.4° radius) appeared in the screen corners (each 12° horizontally and 7° vertically from the screen center). Subjects indicated the perceived expression category via a saccade to the corresponding target (detection window 6° x 6°) (Fig. 1). Target-category associations were learned through reward contingencies assigned by the experimenter. Eye positions outside the bounds of the video or failure to saccade to any of the targets after 10 s aborted the trial. During the training phase, subjects were only rewarded for choosing the correct target. Task training started with expression categories separately in blocks with a single target and a subset of stimuli, then successively more stimuli and more targets were added and more categories were nested until full randomization of trials was reached. The training procedure was flexibly adapted to then animal’s current needs and abilities. Once performance stably exceeded 70 % per category (Supplementary Fig. S1A-B), the test phase was started with new videos presented interleaved with the trained videos in 5-10 % catch trials, in which choice of any target was rewarded. The performance on the trained videos throughout the test phase is shown in Supplementary Fig. S1C-D. In part I of the test phase, all new real macaque videos and the monkey avatar with prototypical expressions were tested. Then, the subjects were specifically re-trained on the prototypical expression avatar videos for approximately two weeks, which were subsequently incorporated into the training set. In part II, the morphed avatar expressions, the scrambled-frame avatar, the point-light avatar, the degraded realism avatars and the human avatar were tested. The morphed test stimuli were presented in 9.9 % of avatars trials (among the prototypical avatar expressions), nested among the original training videos, i.e. in 7.7 % of all trials. Testing sessions were conducted in blocks of 234 to 344 trials (depending of the percentage of catch-trials), with breaks between blocks, and lasted for as long as the animal was willing to work. When the animal stopped working before completion of a block, the remainder of the block was finished in the next testing session. Typically, monkey 1 completed between 1000 and 1500 trials per day and monkey 2 completed between 300 and 900 trials per day. We aimed to collect at least 50 trials per individual test video from each subject. Unfortunately, data collection for part II from monkey 2 could not be completed due to administrative delays in the renewal of our animal testing license. Thus, only 18 (morphed avatar expressions), 30 (scrambled-frame avatar), 29 (point-light avatar), 15 (degraded realism avatars) and 29 (human avatar) repetitions per test video, are available from monkey 2. The human facial expressions, displayed by both the human avatar and the monkey avatar, could only be tested with monkey 1 for the same reason.

**Fig. 1.**
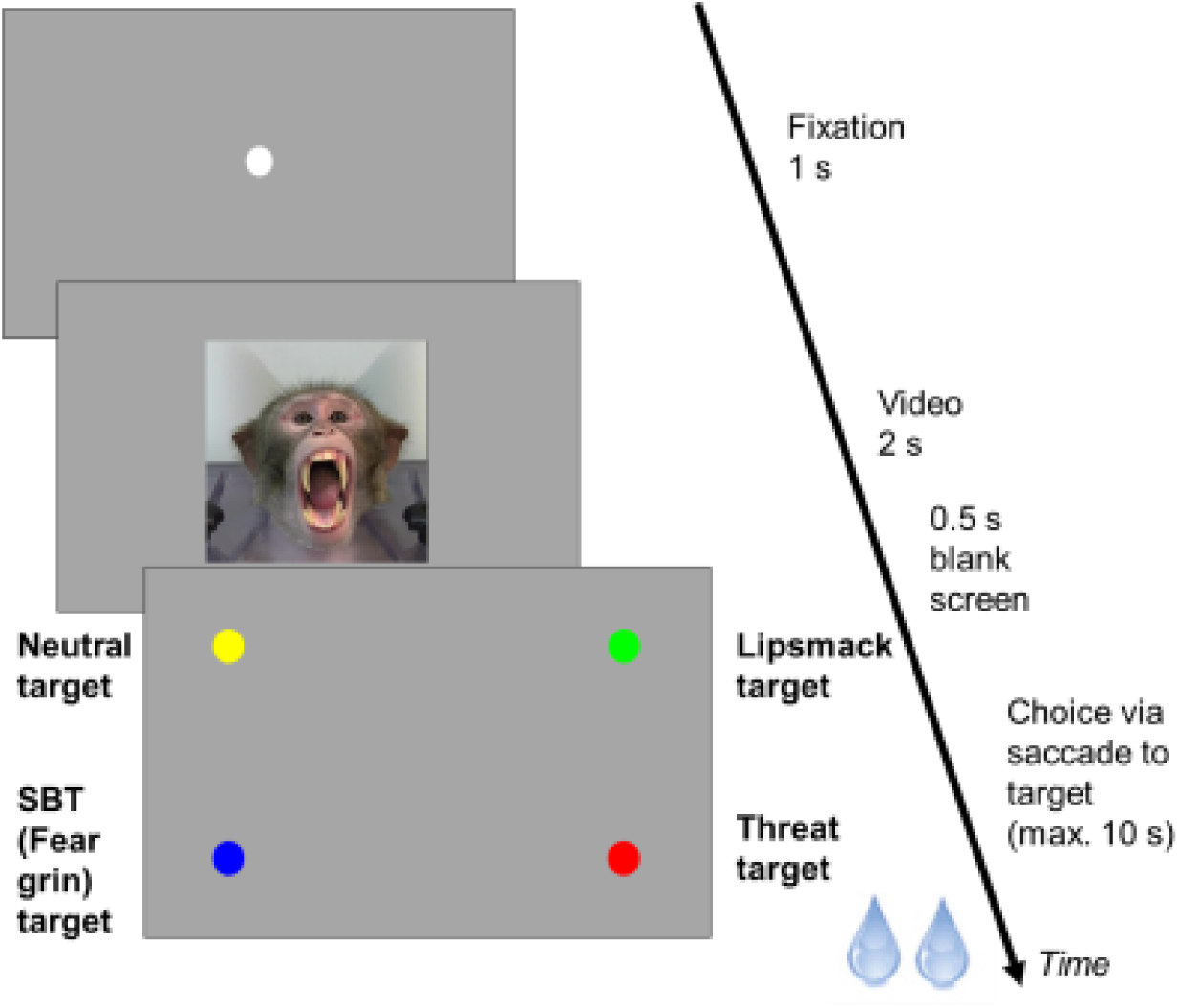
Forced-choice expression categorization paradigm

### Quantification of facial movements

#### Manual FACS coding

MaqFACS AU activations per video were coded before data collection by R. Siebert, who is a certified MaqFACS coder.

#### Automatic FACS coding

Facial movements in the real macaque test videos were tracked via markerless pose estimation implemented in DeepLabCut (DLC) (Mathis et al., 2018; Nath et al., 2019), using the pre-trained “primate face” from the DeepLabCut Animal Zoo, provided by Claire Witham, Centre for Macaques, MRC Harwell, UK (Fig. 2 A) and a model confidence threshold of 0.6. The Eyes_Mid marker was used as a reference and its position was subtracted from all other markers in order to obtain local motion only. For the avatars, mesh joints equivalent to the DLC markers were used (Fig. 2B). AU activations were coded based on the movements of the marker estimates provided by DLC and the avatar mesh coordinates (“automatic coding”). This allowed the metric calculation of AU activation intensity in terms of Euclidian distance from the AU-associated marker position at its neutral starting frame to its position at expression apex (Fig. 2C and detailed in Supplementary Table S1). The thus obtained automatic coding was compared to the initial manual coding by normalizing the scores for each automatically quantified AU between 0 and 1 and subsequently setting all values < 0.1 to 0 (no AU activation) and all other values to 1, thereby mapping the automatic coding onto the format of the manual coding. The agreement between automatic and manual coding could then be compared in the same way as the agreement between two human coders: 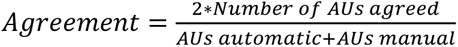

**Fig. 2.**
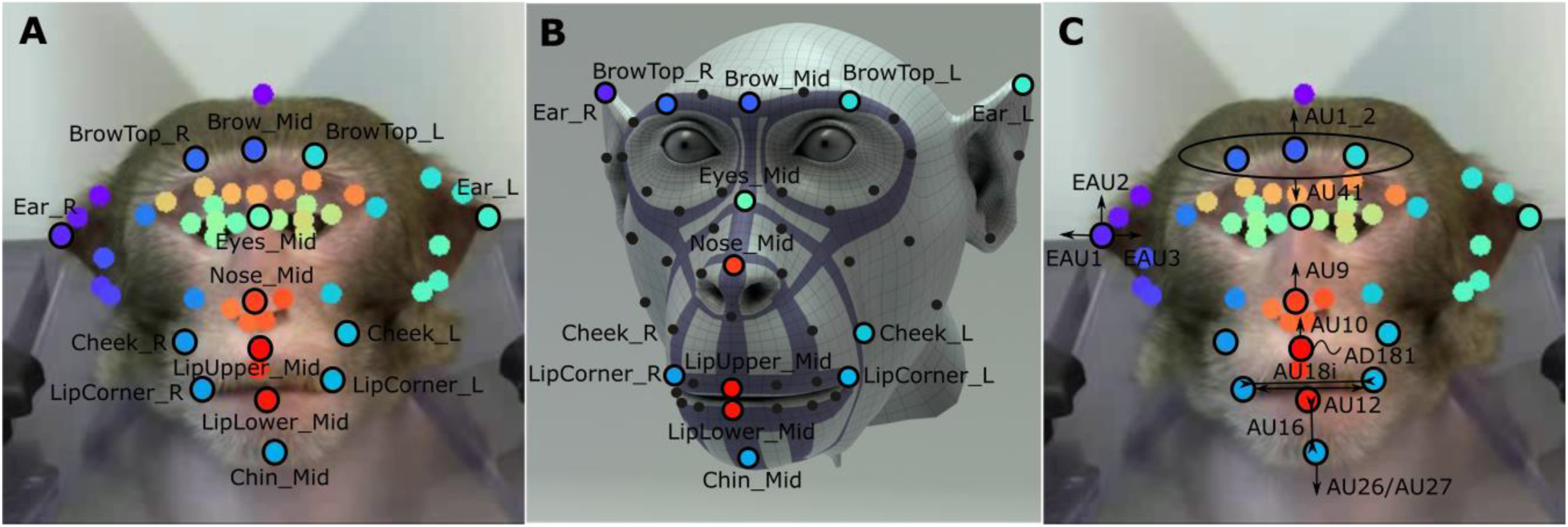
**(A)** Facial movements in the real monkey videos were tracked using DeepLabCut (pre-trained “primate face” model). Markers used to approximate MaqFACS AUs are outlined in black and labeled. **(B)** Avatar mesh joints. Joints equivalent to the employed DLC markers are color-coded and labeled accordingly. **(C)** Schematic of the automatic MaqFACS AU activation and intensity calculation, for details see Supplementary Table S1

The agreement between DLC-tracked real videos and manual coding was 0.72 and the agreement between avatar mesh coordinate trajectories and manual coding was 0.84. Using the minimum agreement of novice coders with certified coders of > 0.7, required to become a new certified MaqFACS coder, we considered the performance of the automatic coding acceptable. Therefore, the quantified automatic coding results were used for all subsequent analyses.

#### Visual similarity

Visual similarities among the different expressions of each real test identity and the naturalistic avatar with prototypical expressions were calculated as the pairwise Pearson correlations between the quantified FACS compositions of the videos (analogous to Micheletta et al., 2015).

### Data analysis

All data were analyzed using MATLAB R2024a and the MATLAB Statistics and Machine Learning Toolbox (MathWorks).

#### Generalization tests

The binomial cumulative distribution function was used to determine whether the relative choice frequency of the category-associated target for each test stimulus exceeded chance (0.25; one-sided *p* < 0.05). We grouped tests into families corresponding to distinct sub-questions: (1) generalization to novel real monkey videos, (2) prototypical avatar expressions, (3) degraded avatar variants, and (4) human expressions. Each family except family 4, which only contained data from one monkey, was additionally split into a) tests over the pooled data of both monkeys and b) tests for both monkeys separately. Within family 2a and 2b, we applied a Bonferroni correction given the small number of comparisons. Within all other families, we applied a Benjamini-Hochberg false discovery rate (FDR) correction, given the larger number of tests and the more exploratory nature of these comparisons.

#### Difference in expected choices according to visual similarity

Expected choice frequencies were equated to the visual similarity matrix (correlations between FACS compositions) and the difference in expected choices was calculated by subtracting the visual similarity matrix from the observed relative choice frequency matrix. The overall Pearson correlation between the visual similarities and the choice frequencies for incorrect targets was calculated (diagonals of visual similarity autocorrelation and correctly chosen targets were excluded).

#### Multinomial Logistic Regression (MLR)

To identify factors driving categorization, we fitted MLR models to the trial-level choices using the MATLAB *fitmnr* function. Predictors included: “Expression” (categorical) for unambiguous expressions or expression intensities of “Lipsmack”, “SBT” and “Threat” (i.e. morphing weights in the range of 0 to 0.75) for the ambiguous morphed expressions, stimulus monkey identity (“Identity”, categorical), stimulus monkey body weight (“Weight”), stimulus monkey averting the gaze away from the camera (“Avert”, scored manually: 0 = direct gaze, 1 = gaze aversion), subject monkey (“Subject”, categorical), and the quantified MaqFACS AU composition of each video were considered as predictors. To avoid multicollinearity among AUs, we omitted redundant AUs (AU8, AU9, AU25) and combined correlated AUs into the following composite predictors: “BrowRaise” (AU1_2), “BrowLow” (AU41), “TeethBare” (AU10+AU12+AU16), “LipPucker” (AU18i*(AD181+1)), “JawOpen” (AU26+AU27), “EarsErect” (EAU1+EAU2) and “EarsFlat” (EAU3). All numeric predictors were z-scored. Choice of the Neutral target was set as reference category and the log-odds of choosing each of the other categories relative to this reference were calculated. Models with different sets of predictors were compared based on the Deviance *D* (generalized residual sum of squares), which measures the goodness of fit compared to a saturated/perfect model (lower values indicating better fit): *D* = −2(log *L*(*b*1, *y*) − log(*bs*, *y*)), whereby *L*(*b*,*y*) represents the maximum value of the likelihood function for a model with the parameters *b. b1* contains the estimated parameters for model 1 and *bs* the estimated parameters for the saturated model. When model 1 has a subset of the predictors in model 2, *ΔD* = *D*2 − *D*1 has a Chi^2^-distribution with degrees of freedom equal to the difference in the number of predictors in model 1 and model 2, allowing for the calculation of a *p*-value. If the *p* < 0.05, we considered model 2 significantly better than model 1 and included the additional predictor(s).

#### Random Forest Classification

To clarify which (potentially non-linear) combinations of AU activation thresholds would lead to which categorization choice, we trained random forest classifiers (500 trees) on the trial-level target choice data using the MATLAB *TreeBagger* function, considering the composite AU predictor set (see MLR) and the full set of AU predictors. Random forests are a robust, non-parametric ensemble learning method (Breiman, 2001) increasingly used for classification problems in animal behavior research (Bergen et al., 2023), and have previously been applied to human facial AU data to classify facial expressions (Pu, Fan, Chen, Ji, & Zhou, 2015) or predict valence and arousal ratings (Zhang et al., 2024). Random forests aggregate predictions from multiple decision trees, each trained on a bootstrapped data sample with random feature subsets considered at each split. The out-of-bag (OOB) error is computed from predictions on data not included in each tree’s bootstrap sample, providing an internal validation metric. Feature importance was quantified using OOB permutation importance scores, which measure the increase in prediction error when each predictor’s values are randomly permuted. Decision rules were extracted through simplified decision trees *(fitctree)* and partial dependence plots, showing how category probabilities change as a function of individual predictor activation while holding the other predictors at their mean values.

#### Pupil size

Raw pupil size (vertical diameter) was preprocessed by first removing blinks (arbitrary eye tracker units < -4) and other artifacts larger or smaller than the mean +/- 2*standard deviation, then smoothing the signal with a 2^nd^ order Savitzky-Golay filter of window size 100 ms and finally translating the signal within each block to *z*-scores. Average pupil size per trial from 1750 ms (when the pupillary light reflex to the movie onset was assumed to have settled) to 2750 ms was calculated, which was the unit for all statistical comparisons. The brightness of each test video frame was calculated as the mean pixel intensity of the grayscale image. The brightness between expressions was neither significantly different for the complete real expression set (Kruskal-Wallis ANOVA, *p* = 0.92) nor the incomplete real expression set missing the SBT (Kruskal-Wallis ANOVA, *p* = 0.69). All avatar videos had been rendered using the exact same lightning and scene settings for each expression. Pupil sizes between expression categories were compared using ANOVAs and *post-hoc* Tukey-Kramer tests. Post-hoc comparisons were based on estimated marginal means from the constrained (Type III) ANOVA model, which adjusts for unequal trial numbers across subjects and identities rather than weighting by raw cell sizes. General Linear Models (GLMs) were fitted to the pupil size data using the same stepwise approach and the same set of predictors as for the MLR models of the choice behavior, plus the additional predictor video brightness and the control predictors total AUs intensity (operationalized as the sum of all normalized (range 0, 1) AU scores per video) and number of activated AUs (operationalized as the number of AUs with normalized scores >= 0.1 per video, analogous to the operational definition of AU activation for the automatic to manual coding comparison).

## RESULTS

### Part I: Naturalistic monkey stimuli

#### Partial generalization of facial expression categorization to new videos with differential pupil reaction to the expression categories

After the monkeys had been trained to categorize videos of conspecifics showing neutral, lip-smacking, silent bared-teeth (SBT) and open-mouth threat facial expressions, they could to some extent generalize the learnt categories to new monkey identities, including a naturalistic monkey avatar. The threat expression was largely correctly categorized and also the lip-smacking expression was often reliably categorized as such, however rarely distinguished from neutral expressions, whereas the classification of the SBT display varied the most (Fig. 3A). Overall, the correct classification of real expressions was moderate with 42.01 % (neutral 20.96 %, lip-smacking 53.04 %, SBT 26.08 %, threat 67.97 %). Similarly, the correct classification of the monkey avatar videos was 38.56 % (neutral 48.21 %, lip-smacking 27.68 %, SBT 13.16 %, threat 65.18 %) (Fig. 3C). A Multinomial Logistic Regression (MLR) analysis of the real monkey video and naturalistic avatar categorization choices with “Expression” as categorical predictor corroborated that the expression category significantly predicted the target choice (MLR Model 1: *Chi^2^*(9, 4086) = 1593.55, *p* < 0.001; Fig. 4B and Supplementary Table S2).

**Fig. 3.**
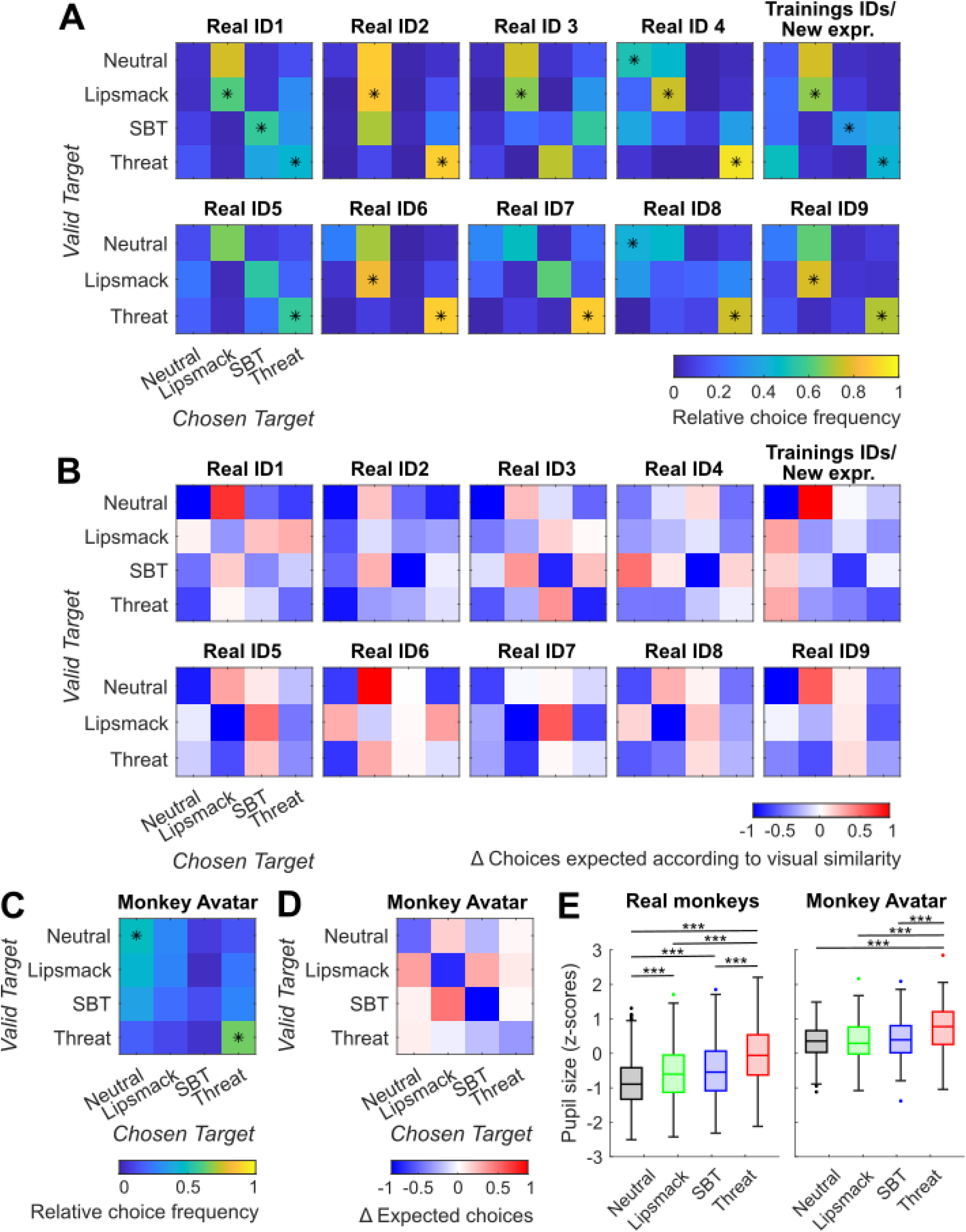
**(A)** Confusion matrices of categorization frequencies for all real monkey test identities, averaged performance of the two monkeys. Asterisks indicate significant generalization (binomial tests, chance level 0.25, *p* < 0.05, FDR-corrected). **(B)** Difference matrices for all real monkey test identities, averaged performance of the two monkeys. Differences were calculated between observed relative choice frequency and the expected choice frequency according to the FACS-based visual similarity between the test expression and the expression associated with the selected target. A value of zero (white) indicates that the target was selected as often as predicted by the visual similarity, positive values (red) indicate that the target was selected more often, and negative values (blue) indicate that the target was selected less often. **(C)** Confusion matrix of categorization frequencies for the naturalistic monkey avatar with prototypical expressions, averaged performance of the two monkeys. Asterisks indicate significant generalization (binomial tests, chance level 0.25, *p* < 0.05, Bonferroni-corrected). **(D)** Difference matrix for the naturalistic monkey avatar with prototypical expressions. **(E)** Boxplots of pupil sizes (both monkeys) during the viewing of each expression category for all real monkey test identity expressions (left) and the monkey avatar with prototypical expressions (right). \*\*\**p* < 0.001 (ANOVAs, *post-hoc* Tukey-Kramer tests). Systematic pupil size differences between real monkey videos and monkey avatar are influenced by differences in stimulus background brightness

**Fig. 4.**
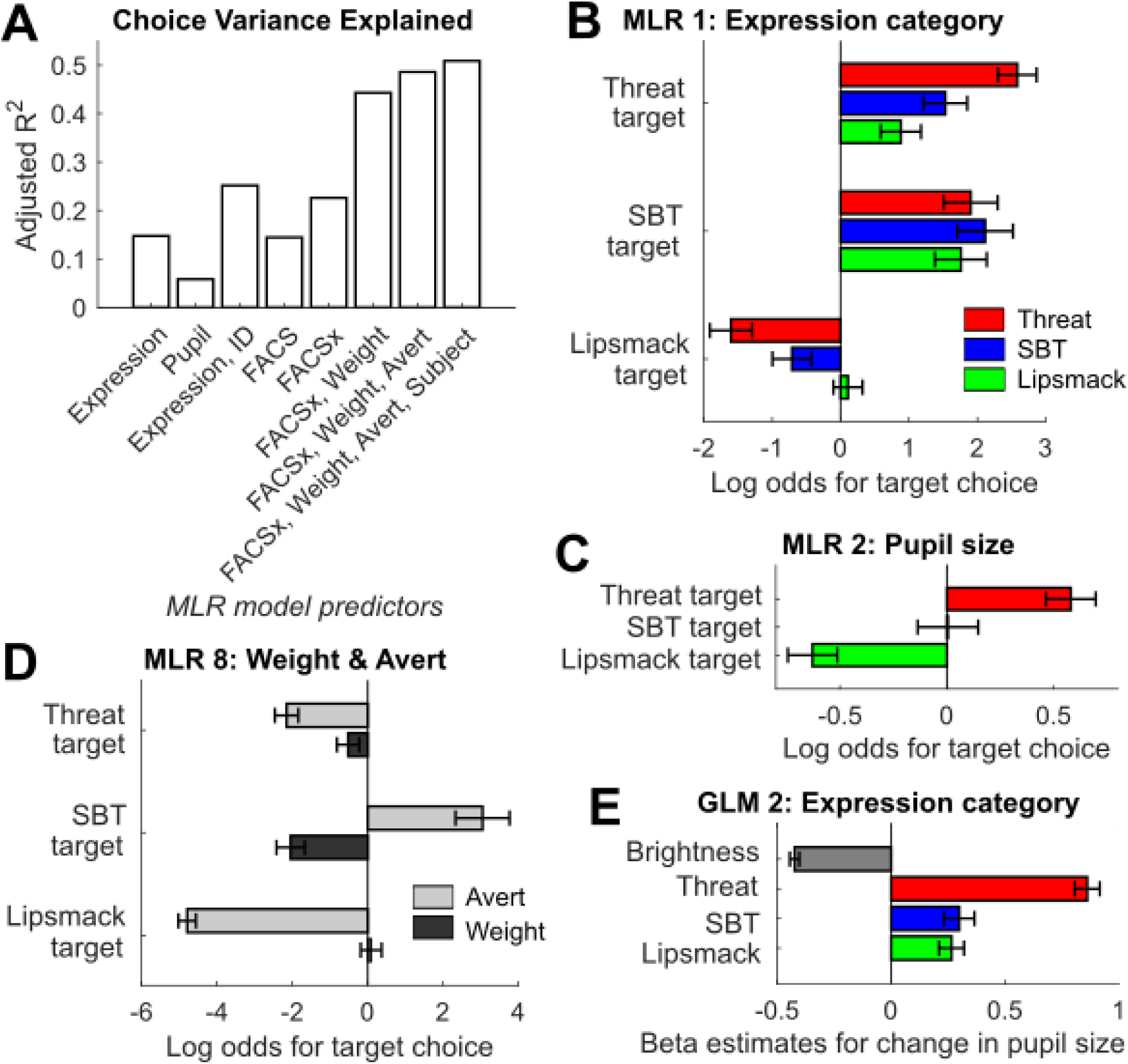
Models of choice and pupil size data in response to the pure (real and naturalistic monkey avatar) expression videos. All model details are available in Supplementary Table S2 (choice) and Supplementary Table S4 (pupil size). Error bars represent 95% confidence intervals. **(A)** Proportion of the variance in choice behavior explained (adjusted R^2^) by MLR models with different sets of predictors. “FACS” stands for the composite FACS predictors BrowRaise (AU1_2), BrowLow (AU41), TeethBare (AU10+AU12+AU16), LipPucker (AU18i*(AD181+1)), JawOpen (AU26+AU27), EarsErect (EAU1+EAU2) and EarsFlat (EAU3). “FACSx” stands for the composite FACS predictors + 3-way interactions between mouth predictors (LipPucker, TeethBare. JawOpen) x brow predictors (BrowRaise, BrowLow) x ear predictors (EarsErect, EarsFlat). **(B)** Log odds coefficients for choosing each target relative to choosing the Neutral target (reference category) in MLR model 1 with expression category as predictor. **(C)** Log odds coefficients of MLR model 2 with pupil size as predictor. **(D)** Log odds coefficients for the predictors body weight and averted gaze in MLR model 8; full model results are available in Supplementary Table S3. **(E)** Beta coefficients in GLM Model 2 with expression category and video brightness as predictors of the pupil size

Errors were not explained by FACS-based visual similarity between expressions alone, as the visual similarity, in terms of correlation between the FACS compositions of expression pairs, was not correlated with the choice frequency of an incorrect target, neither for both monkeys together (Pearson’s *rho* -0.079, *p* = 0.39), nor for monkey 1 (Pearson’s *rho* -0.13, *p* = 0.20) or monkey 2 (Pearson’s *rho* -0.053, *p* = 0.59) individually. The differences between observed relative choice frequency of each target in response to each expression video and expected choice frequency according to the visual similarity between the shown expression and the expression associated with the selected target for each test identity are presented in Fig. 3B and D. If the monkeys selected a target as often as predicted by the visual similarity, the difference in expected choices would be zero (white), whereas the frequent positive (red) and negative (blue) values indicate that targets were selected more and less often as visual similarity would predict. Performances and differences in expected choice frequencies for both monkeys separately, along with the visual similarity matrices, are depicted in Supplementary Fig. S2 and Supplementary Fig. S3.

Analysis of the pupil size during the stimulus presentation underlines the facilitated recognition of threat and suggests that the expression categories are perceived differentially despite partial lack of behavioral distinction, as pupil size was significantly different between the seen facial expression categories. Pupil size is a measure of autonomic arousal with larger pupils found in states of higher arousal (Grujic, Polania, & Burdakov, 2024) and has been shown to change in reaction to socially important rhesus macaque stimuli (Ebitz, Pearson, & Platt, 2014). In our study, pupils were overall largest, indicating highest arousal, when seeing threatening expressions, second largest when presented with the SBT or the lip-smacking display, and smallest when the stimulus monkey kept a neutral expression (real monkey videos, one-factorial ANOVA for Expression: *F*(3, 3620) = 211.42, *p* < 0.001), see Fig. 3E left and Supplementary Fig. S4 for the pupil sizes of both monkeys during the viewing of each test stimulus separately. Also for the naturalistic avatar alone, pupils were significantly enlarged when viewing the threatening expression (one-factorial ANOVA for Expression: *F*(3, 446) = 11.80, *p* < 0.001) (Fig. 3E right). A General Linear Model (GLM) analysis of the pupillary reactions to the real monkey and naturalistic avatar expressions confirmed these results while explicitly accounting for the influence of video brightness (GLM Model 2: *F*(4, 4069) = 593.56, *p* < 0.001; Fig. 4E and Supplementary Table S4). Conversely, pupil size alone was significantly predictive of the choice behavior in an MLR: larger pupils predicted choice of the Threat target and smaller pupils predicted choice of the Lipsmack target. (MLR Model 2: *Chi^2^*(3, 4074) = 631.30, *p* < 0.001; Fig 4C and Supplementary Table S2).

#### Interactions between face parts and additional social information influence the categorization behavior

Examination of the categorization performances for each test identity in Fig. 3A and of the individual performances of both monkeys (Supplementary Fig. S2) suggested that the monkeys used idiosyncratic categorization strategies for different test identities. This was confirmed by an MLR analysis with “Identity” added to “Expression” as predictors of the target choice, which significantly improved the model fit (MLR Model 3: *ΔD* = 1184.44, *p* < 0.001; Supplementary Table S2 and Fig. 4A). We therefore wondered what factors exactly in the faces and facial movements of the different test identities might lead to these differences in categorization. We hypothesized that subtle variations in the exact facial movement compositions (eyebrow and ear movements, degree of mouth movements), information about the monkey’s perceived strength, or direct vs. averted gaze might cause facial expressions to be perceived differently, and that the two monkey subjects might have pursued individual categorization strategies. To test these hypotheses, we compared different MLR models in a step-wise procedure, the results of which are summarized in Supplementary Table S2 and Fig. 4A. In order to describe facial movements agnostic of *a priori* category labels, we used the MaqFACS framework to identify and quantify activated facial action units (AUs) and thus deconstruct the facial expressions into their muscle movement components. The composite FACS predictors (MLR Model 4: *D* = 9048.91, *Chi^2^*(21, 4086) = 1584.04, *p* < 0.001) yielded similar explanatory power as using the expression category predictor (MLR Model 1: *D* = 9039.40, *Chi^2^*(9, 4086) = 1593.55, *p* < 0.001). However, including three-way interactions between the mouth predictors (“LipPucker”, “TeethBare”, “JawOpen”), the brow predictors (“BrowRaise”, “BrowLow”) and the ear predictors (“EarsErect”, “EarsFlat”), which we abbreviated as “FACSx”, yielded a significantly improved model (MLR Model 5: *ΔD* = 940.64, *p* < 0.001). Moreover, the monkeys did not seem to judge only the shown expression, but rather interpret a facial display differently depending on the stimulus monkeys’ weight (assumed to be associated with perceived strength), and depending on whether the facial display included averted gaze, as adding the predictors “Weight” (MLR Model 6: *ΔD* = 2311.96, *p* < 0.001) and “Avert” (MLR Model 7: *ΔD* = 461.58, *p* < 0.001) both yielded significantly improved models. Finally, the two subject monkeys’ individual strategies (“Subject”) also had a small, but significant influence (MLR Model 8: *ΔD* = 248.41, *p* < 0.001; see also Supplementary Table S3). Higher body weight significantly decreased the probability that the SBT target would be chosen, whereas averting of the gaze significantly decreased the probabilities for the choice of the Lipsmack and Threat targets, and increased the probability for the choice of the SBT target (Fig. 4D and Supplementary Table S3).

Identical stepwise GLM analysis was performed for the pupil size whilst taking video brightness into account for all models (Supplementary Table S4 and Supplementary Fig. S5). Analogously to the choice behavior, the pupil size was similarly well explained by expression category (GLM Model 2: *D* = 1835.38, *F*(4, 4069) = 593.56, *p* < 0.001) and FACS composition (GLM Model 4: *D* = 1862.84, *F*(8, 4065) = 284.63, *p* < 0.001) and model fit was improved by accounting for interactions between the mouth, brow and ear AU predictors (GLM Model 5: *ΔD* = 107.79, *p* < 0.001), as well as by including “Weight” (GLM Model 6: *ΔD* = 379.76, *p* < 0.001), “Avert” (GLM Model 7: *ΔD* = 58.07, *p* < 0.001) and “Subject” (GLM Model 8: *ΔD* = 123.02, *p* < 0.001) as predictors. Control analyses using the total intensity of AUs in the video (GLM Model 9: *D* = 2044.42, *F*(2, 4071) = 858.13, *p* < 0.001) or the number of activated AUs (GLM Model 10: *D* = 2082.14, *F*(2, 4071) = 805.71, *p* < 0.001) as predictors explained the pupil size markedly worse than the expression category or the FACS composition, indicating that the amount of motion *per se* did not drive arousal.

### Part II: Synthetic stimuli (avatar variants)

#### Categorization of morphed expressions reflects their position in morphing space and pupil size reflects the threat intensity

Having found that facial expression perception is influenced by a number of factors beyond the expression itself, namely depending on who is showing the expression and where exactly the individual is looking, we investigated with a fully controlled avatar stimulus set, 1) which combination of facial features lead to the perception of a facial expression category, and 2) which facial shape and appearance features are necessary and sufficient for a facial movement to be perceived as falling into one of the trained facial expression categories. Before this second test phase, the pure, prototypical facial expression videos of the naturalistic avatar were added to the training stimulus set and the monkeys were briefly trained again until they could reliably categorize the avatar videos. To answer question 1), we generated a stimulus set of morphed expressions (depicting one identity with constant direct gaze), whereby expressions were interpolated continuously between two prototypical facial expressions in steps of 25 % intensity (Fig. 5). The target choice frequencies, presented in Fig. 6 and Supplementary Fig. S6 (performance of monkey 1 and 2 separately) show that the categorization of the morphed avatar videos followed the morph axes, i.e. the stronger a facial expression category was represented in a video, the more frequently the video was classified as belonging to this category. Clearer lip-smacking features, compared to SBT or Threat features, seemed necessary though for an expression to be categorized as Lipsmack, particularly along the Lipsmack – SBT morph axis, and a general preference for the SBT target was apparent.

**Fig. 5.**
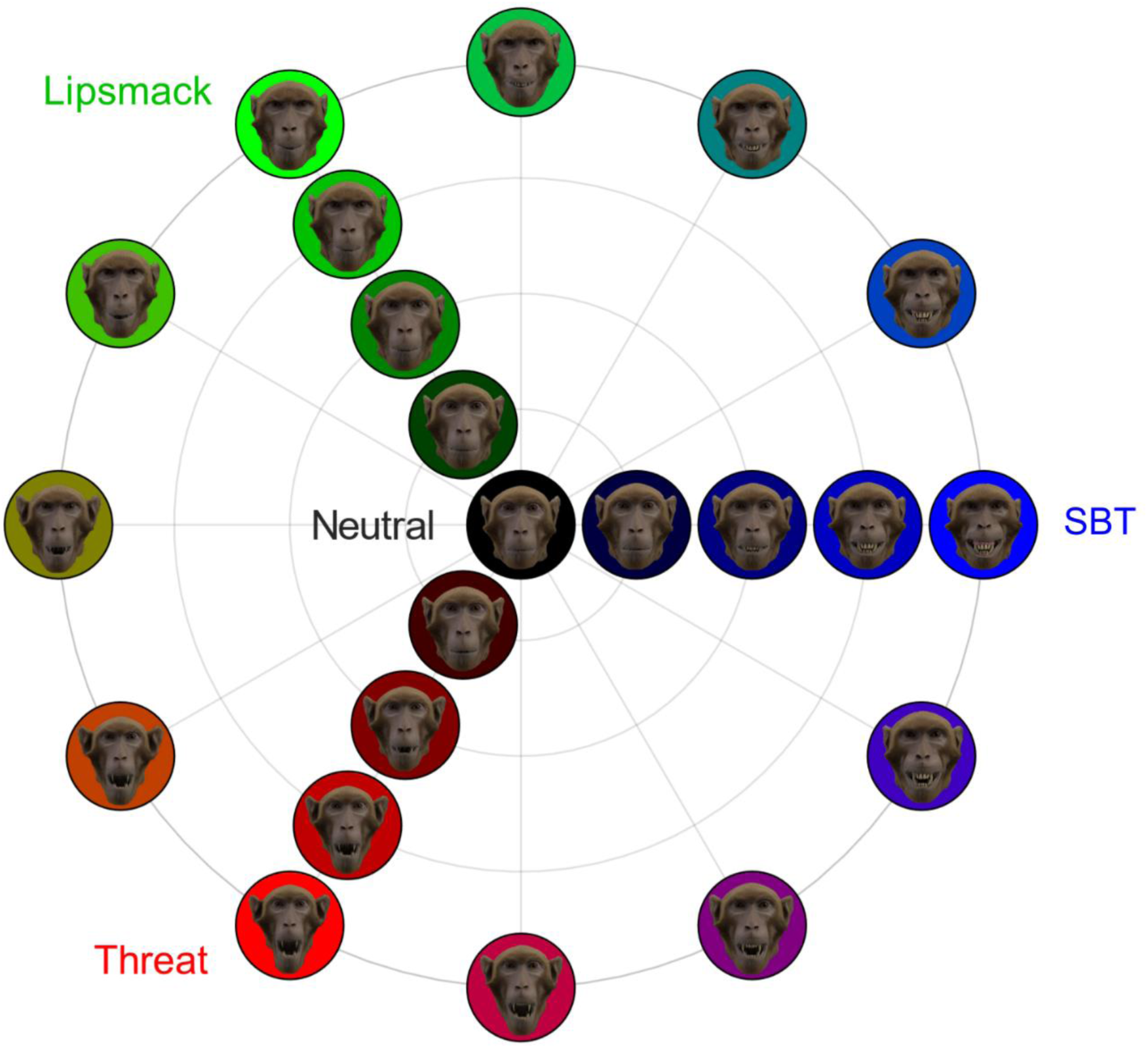
Monkey avatar expression space, comprising the pure, motion capture-based expressions neutral, lip-smacking, silent bared-teeth display (SBT) and open-mouth threat, and graded morphs in steps of 25 % between two pure expressions

**Fig. 6.**
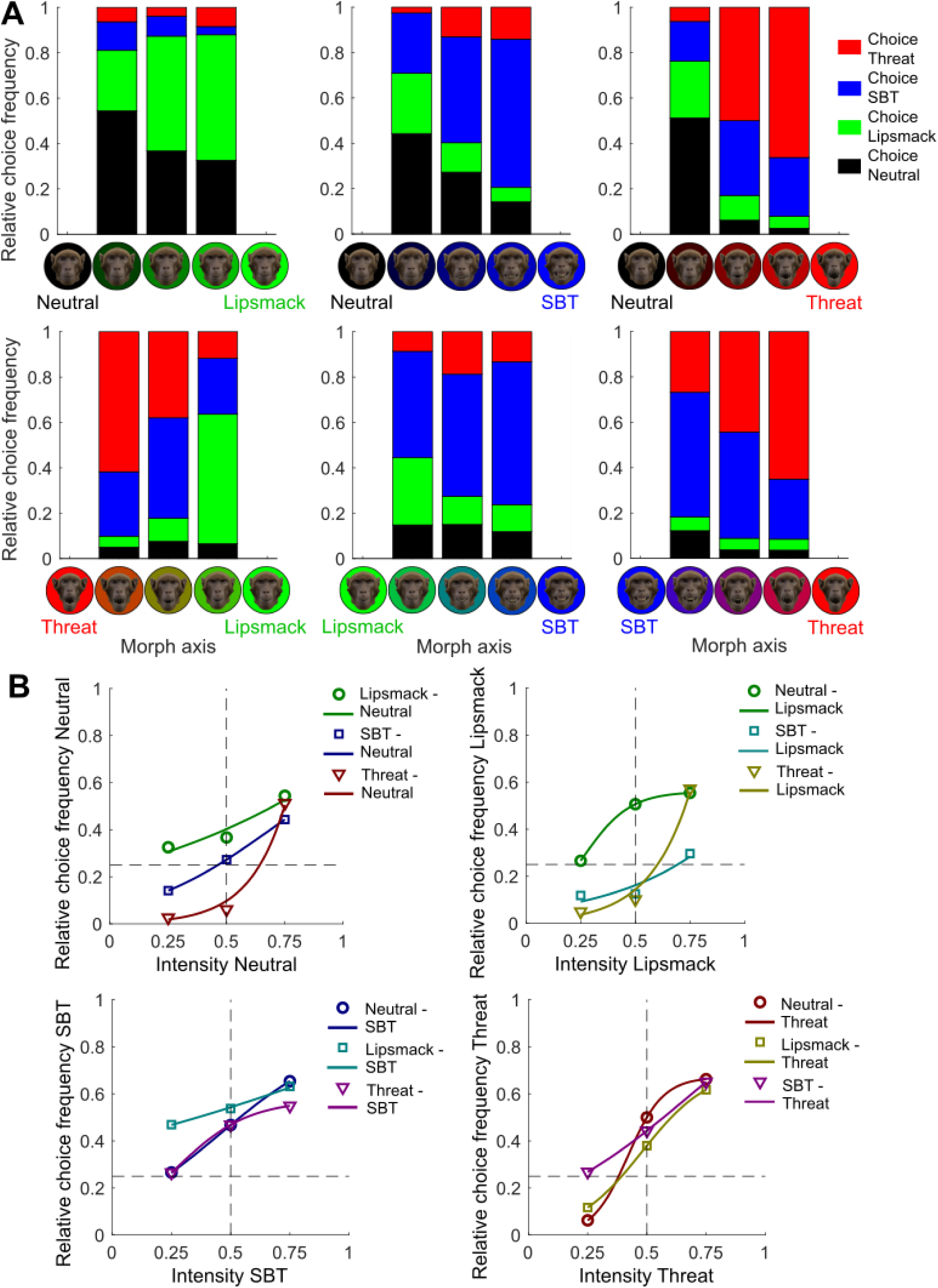
**(A)** Relative choice frequencies for all morphed avatar videos, shown for each morph axis between the prototypical expressions Neutral, Lipsmack, SBT and Threat. **(B)** Relative choice frequencies of the Neutral-, Lipsmack-, SBT- and Threat-associated targets for each morph axis, plotted against the intensity of the associated expression in the morphed video. Horizontal dashed lines indicate the chance choice levels of 0.25, vertical dashed lines indicate the 0.5 expression intensity levels. Logistic fits are overlaid

MLR analysis (Supplementary Table S5) showed that the choice behavior was very similarly well predicted by the intensities of “Lipsmack”, “SBT” and “Threat” in the morphed expression (Morphs MLR Model 1: *D* = 3180.25, *Chi^2^*(9, 1432) = 706.41, *p* < 0.001; Fig. 7A) as by the FACS compositions (Morphs MLR Model 3: *D* = 3164.55, *Chi^2^*(21, 1432) = 734.73, *p* < 0.001; Fig. 7B), which suggests that the category transitions were well captured by the AU metrics. Adding two-way interaction terms to the expression intensities yielded only a small improvement (Morphs MLR Model 2: *ΔD* = 28.33, *p* < 0.001). When attempting to add interaction terms to the FACS AU predictors, analogously to the analysis in part I, maximum likelihood estimation failed to converge because the design matrix was nearly singular, indicating multicollinearity in the predictor matrix, likely resulting from lack of feature variance among the standardized morphed expressions. Adding “Subject” as predictor (Morphs MLR Model 4: *ΔD* = 3.80, *p* = 0.051) did not significantly improve the model, demonstrating that the two monkeys did not adopt diverging strategies to categorize the morphed videos.

**Fig. 7.**
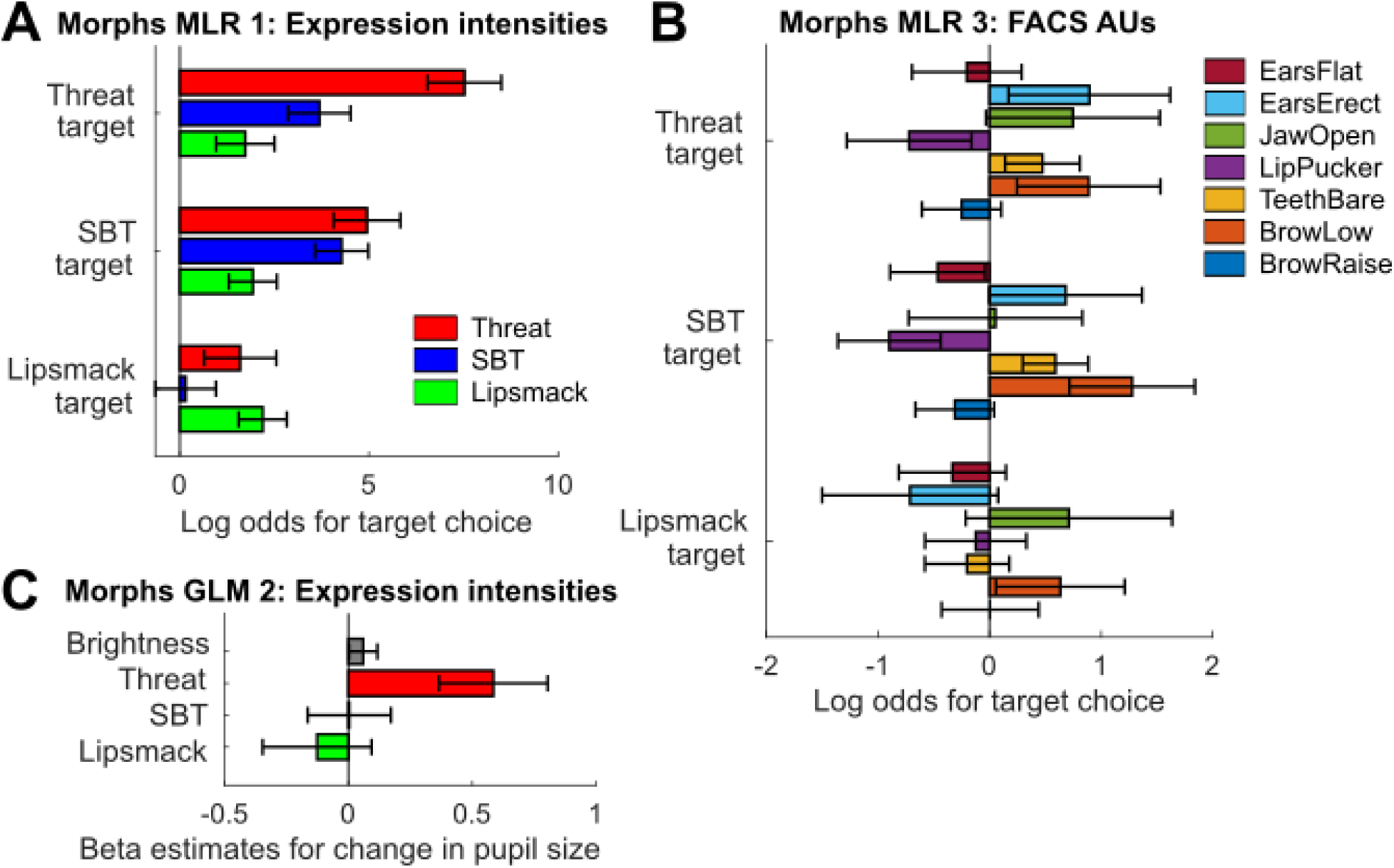
Select models of choice and pupil size data in response to the morphed avatar expressions. All model details are available in Supplementary Table S5 (choice) and Supplementary Table S6 (pupil size). Error bars represent 95% confidence intervals. **(A)** Log odds coefficients for choosing each target relative to choosing the Neutral target (reference category) in Morphs MLR Model 1 with expression intensities in the morphed expressions as predictors. **(B)** Log odds coefficients in Morphs MLR Model 3 with the composite FACS predictors. **(C)** Beta coefficients in Morphs GLM Model 2 with expression component intensities in the morphed expressions and video brightness as predictors of the pupil size

Analogous GLM analysis for the pupil size under inclusion of video brightness as predictor yielded small, but significant effect sizes (Supplementary Table S6). Again, expression component intensities (Morphs GLM Model 2: *D* = 639.19, *F*(4, 1423) = 35.32, *p* < 0.001) explained pupil reactions similarly well as the FACS composition of the video (Morphs GLM Model 4: *D* = 635.15, *F*(8, 1422) = 18.85, *p* < 0.001), but considering interactions among the expression intensities did not significantly improve the model video (Morphs GLM Model 3: *ΔD* = 1.92, *p* = 0.59), whereas adding interaction terms to the FACS AU model again resulted in a rank deficient predictor matrix. Including “Subject” resulted in a marginally better model (Morphs GLM Model 5: *ΔD* = 10.59, *p* = 0.001). As evident from Morphs GLM Model 2 in Fig. 7C and from Supplementary Fig. S7, pupil size was larger the higher the intensity of threat components in an expression, while Lipsmack and SBT component intensity in the morphed expressions did not consistently influence pupil size to one direction. Control analyses using the total intensity of AUs in the video (Morphs GLM Model 6: *D* = 677.87, *F*(2, 1428) = 23.78, *p* < 0.001) or the number of activated AUs (Morphs GLM Model 7: *D* = 679.06, *F*(2, 1428) = 24.66, *p* < 0.001) as predictors explained the pupil size not or only marginally better than the brightness alone and markedly worse than the expression intensities or the FACS composition, which shows that motion intensity *per se* did not increase arousal.

#### Eyebrow and ear movements influence the categorization of ambiguous expressions

To further elucidate which facial movement features decided categorization choices at which threshold, we trained Random Forest Classifiers (RFC), which are based on decision trees and thus naturally handle multicollinearity, complex interactions and non-linear relationships among predictors without the need for explicit modeling. The first RFC used the same set of composite FACS predictors as for the MLR analysis (composite model). Classification accuracy of 53.6 % (OOB error 0.464) exceeded chance level (25 %) and was 18.8 % better than a baseline model only predicting the most frequently chosen category (the SBT target). This modest prediction accuracy is expected given the ambiguous nature of morphed expressions near category boundaries. Classification results are shown in Fig. 8A.

**Fig. 8.**
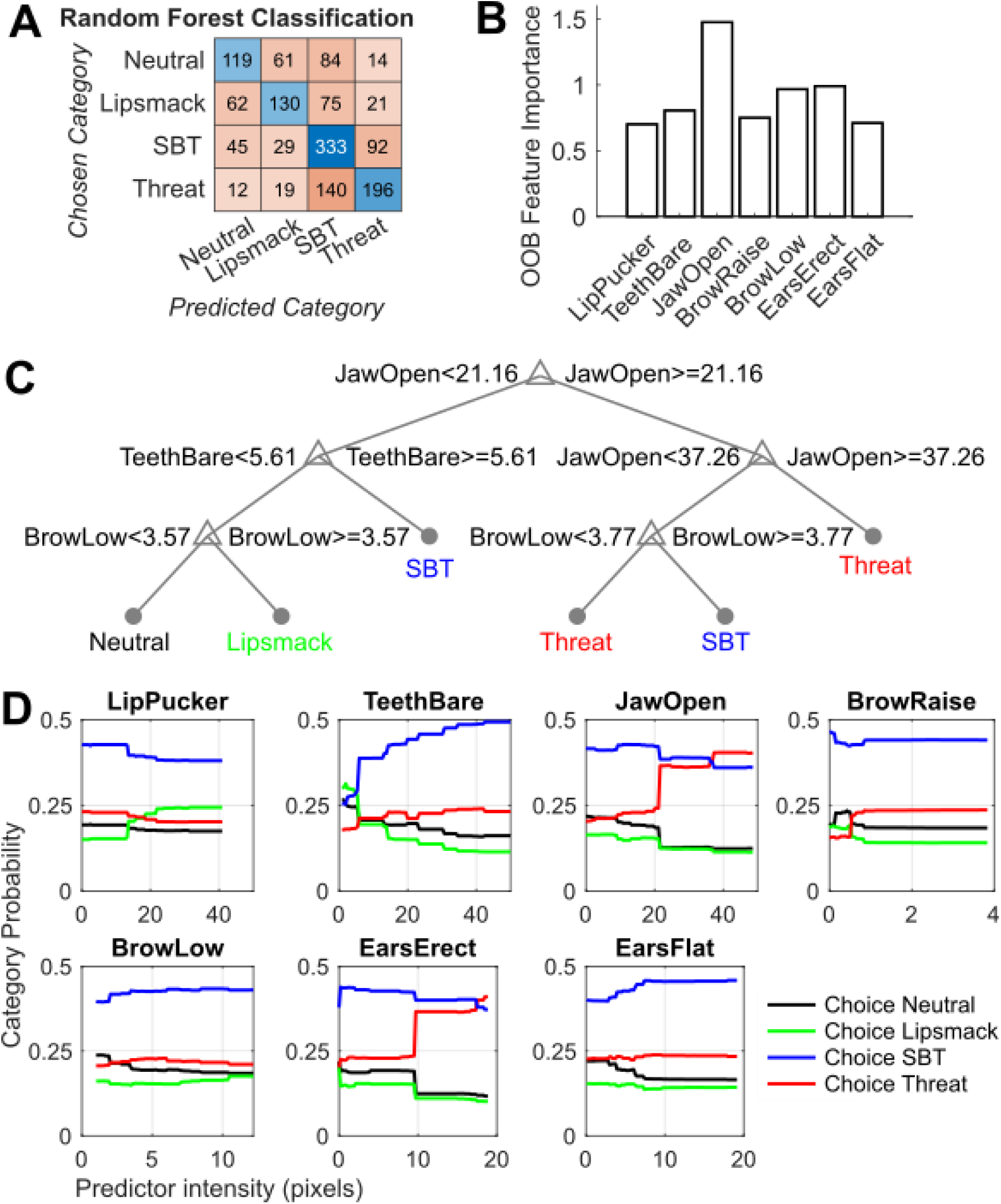
Random Forest Classification results of the morphed avatar expressions using the composite FACS predictors “BrowRaise”, “BrowLow”, “TeethBare”, “LipPucker”, “JawOpen”, “EarsErect” and “EarsFlat” (composite model). **(A)** Heatmap showing the predicted category vs. the chosen category. **(B)** Out-of-bag feature importance of the predictors. **(C)** Simplified decision tree elucidating which combinations of predictor activation intensities lead to which category choices. **(D)** Partial dependency plots showing how individual predictor intensities influence the probability for a category to be chosen

Given the robustness of tree-based methods against multicollinearity – correlated predictors simply compete for inclusion rather than inflating each other’s apparent importance, leaving predictions unbiased (Breiman, 2001) – we asked whether the full set of AUs might yield better prediction and compared the composite model to a model using the full set of individual AU predictors (full AU model). We obtained comparable prediction accuracy with the full AU model (OOB error 0.457, Supplementary Fig. S8A), which is why we focus on the results from the more parsimonious composite model, which also facilitated comparison with the MLR findings.

Our analysis revealed JawOpen as the most important feature (Fig. 8B), which was also the most frequent root split in 35.0 % of trees, while TeethBare was used most frequently overall in 18.2 % of splits. Similarly, the most important feature in the full AU model was AU25 (lips parted) (Supplementary Fig. S8B), used in 11.0 % of splits and as most frequent root split in 30 % of trees, which makes sense as parted lips are involved both in teeth baring and jaw opening, but limits the diagnostic value of AU25 for distinguishing between SBT and threat expressions, which is why we omitted it from the set of composite predictors. A simplified decision tree (accuracy 54.3 %) shows that given large, but ambiguous degrees of jaw opening, lowered brows led to the selection of the SBT target over the Threat target, and given small degrees of both jaw opening and teeth baring, lowered brows decided the choice of the Lipsmack target instead of the Neutral target in our morphed stimulus set (Fig. 8C). This confirms MLR results indicating that brow and ear predictors also significantly influence category choices in addition to mouth predictors (Morphs MLR Model 3, Fig. 7B). A simplified decision tree for the full AU model prompts similar conclusions (Supplementary Fig. S8C). Partial dependency plots elucidate how category choice probabilities change as a function of a single predictor value. Sharp category probability transitions occurred expectedly in TeethBare (max. probability change 24 % towards the SBT target), JawOpen (max. probability change 20 % towards the Threat target) and LipPucker (max. probability change 10 % towards the Lipsmack target), but also in EarsErect (max. probability change 21 % towards the Threat target) or BrowRaise (max. probability change 8 % towards the Threat target) (Fig. 8D).

In order to compare the observed feature importance in the morphed expressions to the unambiguous expressions, we also fitted the composite model to the categorization choices for all pure expression videos (real monkey videos and naturalistic avatar). Trained with the categorizations of these unambiguous expressions, model accuracy increased to 64.5 % (Supplementary Fig. S9A), 26.8 % better than a baseline model predicting only the most common category in this data set (the Lipsmack target). As for the morphed expressions, JawOpen was the most important feature (Supplementary Fig. S9B) and comparable probability changes depending on individual predictor values were observed (Supplementary Fig. S9D). Examination of a simplified decision tree suggests that for unambiguous expressions compared to morphed expressions, LipPucker was more often used as a diagnostic feature to identify Lipsmack expressions, while lowered brows still helped the distinction between Neutral and Lipsmack. However, eyebrow and ear movements seemed overall less influential when the mouth movement was clear (Supplementary Fig. S9C).

A control analysis using an RFC model with the relative intensities of Lipsmack, SBT and Threat in the morphed expressions as predictors yielded nearly identical prediction accuracy as the composite model (OOB error 0.460, Supplementary Fig. S10A) and ascertained Threat intensity as the most important feature (Supplementary Fig. S10B) and most frequent root split (67.6 % of trees). A simplified decision tree for this model (Supplementary Fig. S10C) suggests that the degree of jaw opening can be equated to Threat intensity, and the degree of teeth baring can be equated to SBT intensity, while obscuring the fact that eyebrow lowering seems to be more influential than lip puckering for deciding on the Lipsmack category within our morphed expression stimulus set. Partial dependency plots (Supplementary Fig. S10D) corroborate what choice frequencies (Fig. 6) indicated: the largest category probability increases occurred between 0.25 and 0.5 expression intensity, with additional smaller increases between 0.5 and 0.75.

#### Facial expression recognition requires biologically plausible texture, but not fully realistic monkey faces or coherent motion

In order to further dissect which attributes are necessary for reliable facial expression perception, we first tested versions of the prototypical avatar expressions with time-scrambled frame order, thus abolishing temporal continuity of the movement. Next, we manipulated the facial shape and appearance in two ways: On the one hand, we reduced the realism of the avatar in a stepwise manner while keeping the same movement pattern as for the prototypical expressions of the naturalistic avatar. We first removed the fur (furless avatar), then the color (gray avatar), then the texture (wire avatar), and finally all discernable facial shape (form-from-motion point-light avatar). On the other hand, we changed the facial shape and appearance completely by changing the species of the avatar head from monkey to human, while displaying the exact same monkey facial expressions.

A three-way ANOVA of the pupil size with factors Expression, AvatarType and Subject, including all two-way interactions, revealed significant main effects of Expression, *F*(2, 1870) = 14.99, *p* < 0.001, AvatarType, *F*(4, 1870) = 1278.45, *p* < 0.001, and Subject, *F*(1, 1870) = 144.67, *p* < 0.001. The Expression x AvatarType interaction was significant, *F*(14, 1870) = 5.70, *p* < 0.001, as was the Subject x AvatarType interaction, *F*(5, 1870) = 19.31, *p* < 0.001. The Subject x Expression interaction was not significant, *F*(3, 1870) = 0.39, *p* = 0.76, showing that the expression effect did not differ between the two monkeys. Given the significant Expression x Identity interaction and the fact that the effect of AvatarType cannot be disentangled from systematic differences in brightness between the different avatars, we conducted post-hoc pairwise comparisons (Tukey-Kramer tests) to investigate expression differences within each avatar type separately.

The scrambled-frame avatars were still well recognized and correctly categorized significantly above chance (Fig. 9C and B, right-most panel), while eliciting pupil reactions comparable to the prototypical monkey avatar and real monkey videos, with arousal increasing from Lipsmack over SBT to Threat (Fig. 9D, right-most panel). While the monkeys categorized the furless and gray avatars reasonably well, struggling only with the Lipsmack expression, the wire avatar without texture, consisting only of the wireframe mesh surface, and the point-light avatar were not categorized, instead the monkeys tended to select only one target for these avatar types (Fig. 9A and B). This behavioral pattern is corroborated by the pupil size, which exhibited expression-dependent modulation for the furless and grayscale avatars (though for the latter only marginally significant), but not for the wireframe avatar or the point-light avatar (Fig. 9D). Note that the (limited) data from monkey 2 suggests that he seemed able to discern the Threat expression of the wireframe avatar, possibly due to the conspicuous downward motion of the jaw for this expression, in the absence of a differential pupil reaction (see Supplementary Fig. S11 for the categorization performances and Supplementary Fig. S12 for the pupil sizes of the two monkeys separately). Moreover, the monkeys seemed to some extent recognize the Neutral, SBT and Threat expressions on the human avatar (Fig. 9A and B) with significant pupil size modulation (Fig. 9D).

**Fig. 9.**
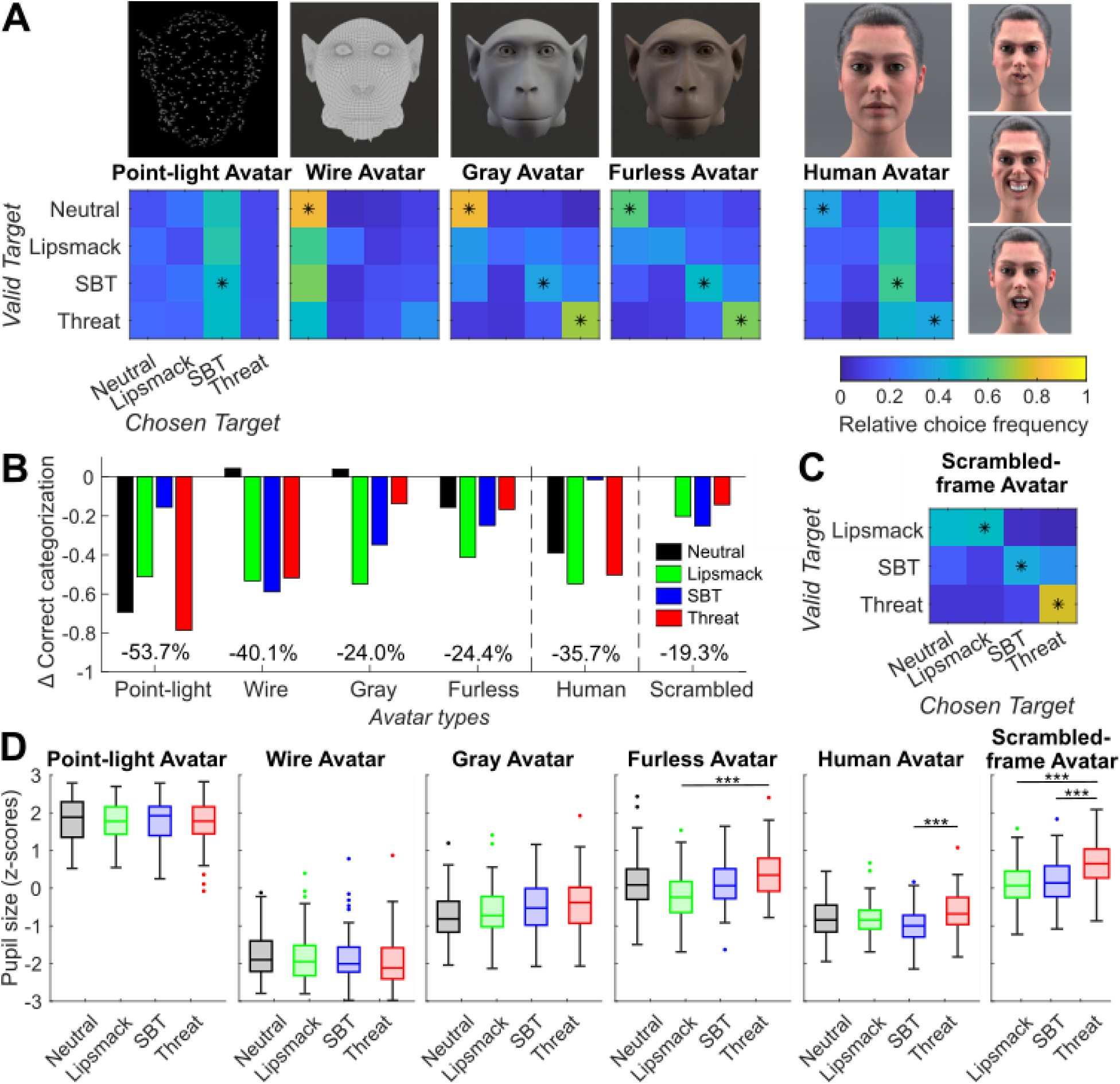
**(A)** Expression categorization frequencies for different avatar variants, asterisks indicate correct categorization significantly above chance level (binomial tests, *p* < 0.05, FDR-corrected). The first four avatar types increase in realism: point-light expressions, wireframe mesh, grayscale avatar and furless avatar. Additionally, a human avatar with monkey expressions was tested; peak lip-smack, SBT and threat expressions (top to bottom) are depicted next to the confusion matrix. **(B)** Difference in fraction of correct categorizations relative to the (now trained) naturalistic avatar for each expression category. Percentages at the bottom indicate the overall difference in correct categorizations. **(C)** Expression categorization frequencies for the naturalistic monkey avatar with time-scrambled frame order (binomial tests, *p* < 0.05, FDR-corrected). **(D)** Boxplots of pupil sizes during the viewing of each expression category for all avatar variants. \**p* < 0.05, \*\*\**p* < 0.001 (ANOVAs, *post-hoc* Tukey-Kramer tests). The difference in pupil size between the Neutral and Threat category for the Gray Avatar marginally missed significance at *p* = 0.082. Systematic pupil size differences between avatar variants are influenced by differences in avatar type brightness

#### Human facial expressions do not fit into the trained categories

We also tested how human facial expressions, displayed by the human avatar or the monkey avatar (Fig. 10A), would be categorized, in order to explore the possibility of a homologous interpretation across primate facial expressions. These stimuli could only be tested with one monkey. We did not find a systematic categorization pattern of the human facial expressions. They were neither categorized according to putative homologous meanings (smiling/happy and lip-smacking, gasping/fearful and SBT, scowling/angry and threat), nor according to potential phylogenetic homology, such as SBT displays and human smiles (Fig. 10B). Moreover, the target choice frequencies were uncorrelated with the visual similarities between the human expressions and the target-associated monkey expressions, measured as the correlation between the FACS compositions of each expression pair (Supplementary Fig. S2A), for the human avatar (Pearson’s *rho* 0.05, *p* = 0.89) and the monkey avatar (Pearson’s *rho* -0.34, *p* = 0.29). The differences between observed relative choice frequency of each target and expected choice frequency according to the visual similarity between the viewed human expression and the target-associated monkey expression are depicted in Fig. 10C. These findings suggest that the human facial expression did not carry meaning for the subject monkey, a notion that is supported by the lack of differential pupil reactions between expressions (Fig. 10D). A two-factorial ANOVA of the pupil size showed neither a main effect of Expression (*F*(2, 381) = 1.53, *p* = 0.22), nor an interaction between Expression and AvatarType (*F*(2, 381) = 1.36, *p* = 0.26), only a significant main effect of AvatarType (*F*(1, 381) = 778.18, *p* < 0.001), likely due to differences in stimulus and background brightness between the human and monkey avatars.

**Fig. 10.**
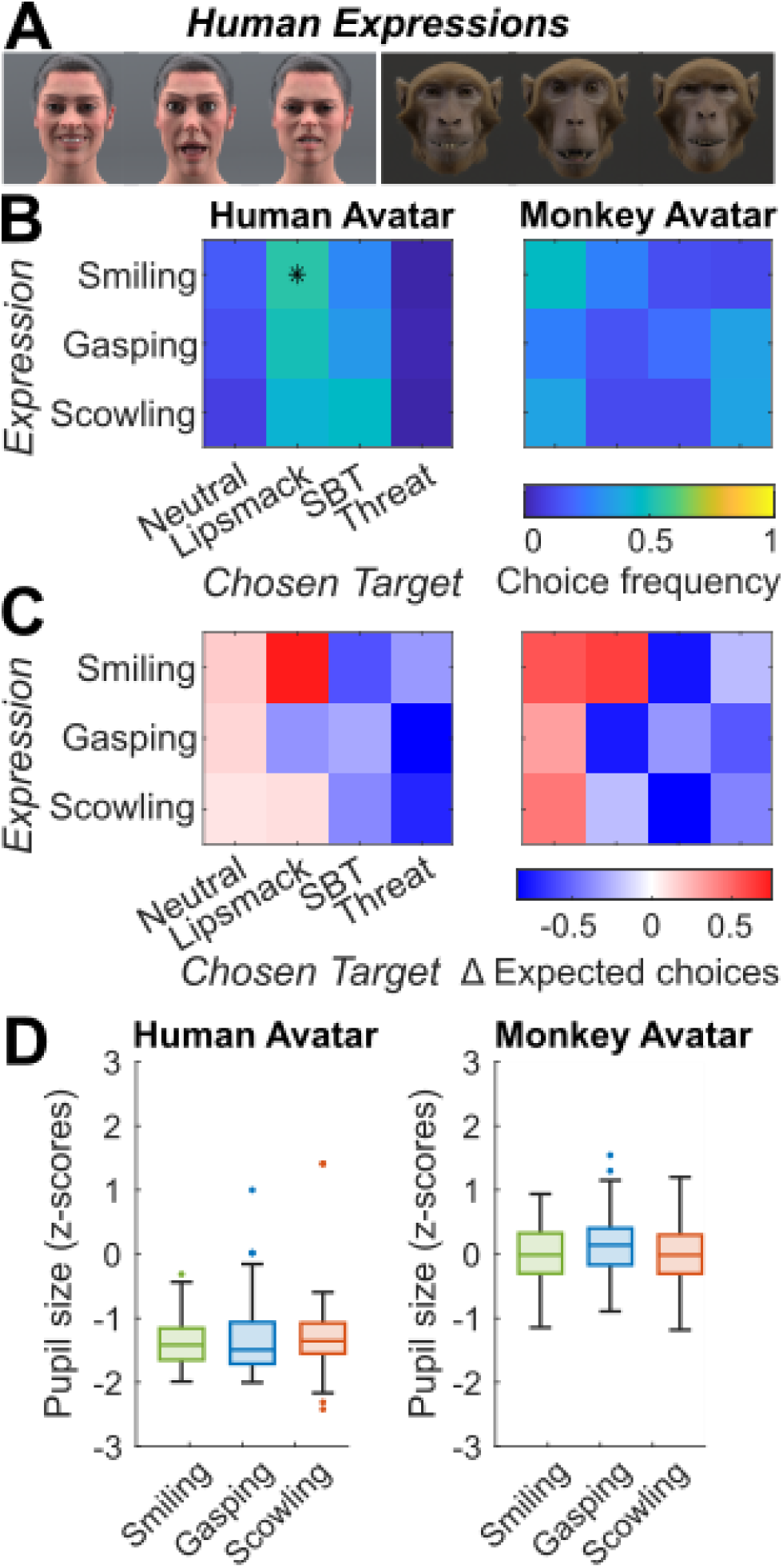
**(A)** Smiling (happy), gasping (fearful) and scowling (angry) human expressions of the human avatar (left) and the monkey avatar (right). **(B)** Expression categorization frequencies of monkey 1 for the human avatar and the monkey avatar displaying the human facial expressions smiling, gasping and scowling (*N* per video = 60). The Lipsmack, SBT and Threat targets, respectively, were considered the “valid” according to putative homologous meanings. Asterisks indicate generalization significantly above chance level (binomial tests, *p* < 0.05, FDR-corrected). **(C)** Difference matrices calculated between observed relative choice frequency and the expected choice frequency according to the visual similarity between the human test expression and the monkey expression associated with the selected target. A value of zero (white) indicates that the target was selected as often as predicted by the visual similarity, positive values (red) indicate that the target was selected more often, and negative values (blue) indicate that the target was selected less often. **(D)** Boxplots of pupil sizes of monkey 1 during the viewing of each human expression category for the human and monkey avatars. Systematic pupil size differences between human avatar and monkey avatar are influenced by differences in stimulus and background brightness, compare (A).

## DISCUSSION

We investigated how rhesus macaques categorize facial expressions, and which facial features enable expression perception. In a forced-choice categorization paradigm with neutral, lip-smacking, silent bared-teeth (SBT) and open-mouth threat displays, two rhesus macaques successfully applied learned categories to unfamiliar individuals and avatars. Pupil size measurements showed graded physiological responses across expression types, demonstrating that the macaques extracted behavioral meaning rather than simply sorting low-level visual patterns. When viewing expressions morphed between two prototypes, categorization frequencies tracked the proportional intensity of each expression component, indicating perception along a continuum. To quantify similarity between expressions, we developed a novel automated approach combining DeepLabCut landmark tracking with the MaqFACS framework, enabling objective, muscle-based quantification of all facial movements in frontal videos of monkey faces without being limited to mouth movements only (Blumrosen, Hawellek, & Pesaran, 2017) or requiring pre-labeled training datasets (Morozov, Parr, Gothard, Paz, & Pryluk, 2021). This approach showed that FACS-based visual similarity between expressions alone did not explain error patterns. Instead, categorization was influenced by how facial regions (mouth, eyebrows, ears) moved in coordination, whether gaze was averted, and by signaler characteristics exemplified by the displaying individual’s body weight. This suggests that the tested expressions types were not interpreted as rigid categories with one meaning, but instead category-defining movements were interpreted in a nuanced manner embedded within their entire facial and signaler contexts. Systematic manipulation of avatar properties showed that neither temporally coherent motion, nor photorealistic rendering or a conspecific face were essential for expression recognition, but discernible facial shape and surface texture were required. Notably, human expressions displayed by human and monkey avatars failed to elicit systematic categorization or differential arousal.

### Categorization of facial expressions does not follow clear morphological categories

In part I, we show that two rhesus macaques trained to categorize facial expressions into neutral, lip-smacking, silent bared-teeth (SBT) and open-mouth threat displays could to some extent generalize these learned categories to new instances of those expressions presented by new agents, both real monkeys and monkey avatars. Critically, viewing these expressions was accompanied by differential arousal as measured by pupil size, with incremental pupillary responses to neutral, lip-smacking, SBT and threatening faces. Addressing our first hypothesis regarding discrimination strategy, the differential physiological arousal precludes that the macaques resorted to the categorization of elementary facial movement features without attaching any meaning to the expressions. If they had relied solely on low-level visual features, we would have expected comparable performance across expression categories without physiological differentiation. Instead, the pattern of arousal responses, which was not explained by motion intensity *per se*, particularly the robust arousal to threatening faces, suggests that these videos were perceived as meaningful categories rather than arbitrary visual patterns. However, the categorization patterns did not align with a perception of the expressions as clear, mouth-movement-defined morphological categories, as miscategorization frequencies differed vastly between expressions, but were not correlated with the visual similarity between expressions as measured by the correlation between their FACS compositions. Open-mouth threat was most frequently recognized as such, reliably differentiated from lip-smacking and elicited the strongest arousal, confirming previous reports of heightened arousal when seeing threatening faces (Dal Monte, Costa, Noble, Murray, & Averbeck, 2015; Landman, Sharma, Sur, & Desimone, 2014; Siebert et al., 2020) and conceivably resulting from an evolutionarily conserved predisposition for prioritized processing of threatening faces (e.g. Frischen, Eastwood, & Smilek, 2008; Gong & Smart, 2021; Kawai, Kubo, Masataka, & Hayakawa, 2016; Öhman, Lundqvist, & Esteves, 2001; Sackett, 1966). Although, note that human data have suggested contesting views (e.g. Becker, Anderson, Mortensen, Neufeld, & Neel, 2011; Nummenmaa & Calvo, 2015; Savage, Lipp, Craig, Becker, & Horstmann, 2013). Lip-smacking, a behavior usually observed in affiliative or appeasing contexts, was correctly categorized for most test monkey identities, but neutral expressions were frequently miscategorized as lip-smacking despite lacking any lip-smacking movement elements, confirming categorization based on factors beyond simple motion patterns. However, the categorization of the SBT display varied substantially across test identities. Using multinomial logistic regression (MLR), we explored which factors might explain these identity-dependent categorization patterns and identified the stimulus monkey body weight, gaze direction (direct vs. averted) and differential interactions between mouth, eyebrow and ear movements as significant predictors within our stimulus set.

After integrating mouth, eyebrow and ear movements, gaze direction and signaler characteristics such as body weight, which criterium could the macaques have used to make their categorization choice? One possibility is that they evaluated the facial displays based on the expressed putative internal states. Ever since Darwin (1872) systematically described facial expressions and proposed that they might relate to inner states, facial behavior in humans has been directly linked to the expression of discrete emotions and widely assumed to be universal (Ekman, 1973, 1992; Ekman & Friesen, 1971; Izard, 1994; Russell, 1994). This notion has only recently been questioned, with growing acknowledgement of the amenability of facial expressions to cultural learning and modulation by individual variation and context (e.g. Barrett, Adolphs, Marsella, Martinez, & Pollak, 2019; Durán & Fernández-Dols, 2021; Le Mau et al., 2021; Valente, Theurel, & Gentaz, 2018). Considering that macaques as nonverbal animals lack semantic emotion concepts, they might sort facial expression images similarly to human dementia patients, who have lost semantic emotion concepts, categorizing them by valence into negative, positive and neutral faces (Lindquist, Gendron, Barrett, & Dickerson, 2014). If the macaques classified the videos according to expressed affect, this would predict a clear separation of positive (presumably lip-smacking) and neutral faces, contrary to our observations. Another possibility is that they sorted the videos according to perceived behavioral intent (Crivelli & Fridlund, 2019; Fridlund, 1994; Frijda & Tcherkassof, 1997; Waller, Whitehouse, & Micheletta, 2017), similarly to a crested macaque (*M. nigra*) that was able to predict social interaction outcomes from facial expressions (Waller, Whitehouse, & Micheletta, 2016). The Relational Model of conflict resolution distinguishes tolerant, avoidant and aggressive intent in social decision making (de Waal & Aureli, 1997). This would explain the lack of differentiation between neutral faces and lip-smacking, as both might convey tolerance of the other, whereas the open-mouth threat display appears to unequivocally convey aggression. The SBT display with exposed teeth and pulled back lip corners has traditionally been described as negatively valued in the majority of anthropoid species, including the rhesus macaque, associated with fleeing, submission or subordination, which can be summarized as avoidant intent (de Waal & Luttrell, 1985; Maestripieri, 1997; Preuschoft, 2000; van Hooff, 1967), and which has brought forth the alternative name “fear-grin”. However, the SBT display is used in more positive contexts in some species, like Tonkean macaques (*M. tonkeana*) (Thierry, Demaria, Preuschoft, & Desportes, 1989), bonobos (Vlaeyen et al., 2022) and humans (the “smile”), where it signals affiliation, appeasement or pleasure, and sometimes dominance, with considerable cultural variation in humans (Rychlowska et al., 2015). Preuschoft and van Hooff (1995) proposed a power asymmetry in the use of this expression across macaque species: While despotic species, such as *M. mulatta*, have a clearer distinction between submission (SBT) and friendliness (lip-smacking), this distinction is blurred in more tolerant species, like *M. tonkeana* or *M. nigra*. For instance, Clark et al. (2020) were able to identify morphological subtypes of the SBT display in *M. nigra* linked to distinct contexts ranging from affiliation over play and copulation to submission. While it is unclear whether such morphological subtypes also exist in *M. mulatta*, context-defined functional subtypes of SBTs have been reported: Conflict SBTs (cSBTs) are shown as signs of immediate submission in aggressive encounters, whereas peaceful SBTs (pSBTs) are preemptively presented as signs of formal hierarchical subordination (Beisner & McCowan, 2014), as previously observed in *M. nemestrina* (Flack & de Waal, 2007). We hypothesize that some SBT displays in our study were interpreted as submission or fear in order to avoid an imminent threat (cSBTs), leading to the choice of the SBT target (or confusion with the Threat target), whereas other SBTs were perceived as more tolerant, appeasing or affiliative (pSBTs), entailing confusion with the lip-smacking category. This interpretation may also explain the inconsistent evaluation of SBTs reported by J. Taubert and Japee (2024). The basis for distinguishing these two SBT subtypes could include subtle facial signs of tension, specific combinations of AUs or AU intensities, or the factors identified by our MLR analysis, body weight and gaze aversion. High body weight of the stimulus monkey reduced the likelihood for the selection of the SBT target, suggesting that an SBT produced by a large, presumably strong monkey was unlikely to be perceived as submission. Direct gaze appeared important for threatening or lip-smacking gestures (associated with approach) to be categorized correctly, as gaze aversion decreased the probabilities for the selection of the Threat and Lipsmack targets. Conversely, gaze aversion (potentially signaling avoidance) increased the chances of an expression to be sorted into the avoidance-congruent SBT category. Averting gaze during a threatening or lip-smacking gesture might create the impression of underlying fear and/or avoidance, contradicting the approach-indicating mouth display, thereby leading with higher probability to the choice of the avoidance-associated SBT category. This aligns with studies in humans reporting a facilitated recognition of approach-associated expressions (happiness and anger) with direct gaze and a facilitated recognition of avoidance-associated expressions (fear and sadness) with averted gaze due to congruent behavioral intent signaled by gaze and facial expression (Adams & Kleck, 2003, 2005). These findings collectively highlight the nuance of primate facial behavior and reveal the limits of mouth movement-defined expression categories, calling for further research into which variations in facial displays carry communicative meaning.

### Mouth, eyebrow and ear movements are integrated to discriminate facial expressions

In part II, we exploited a fully controllable monkey avatar to map out which facial motion and shape features are important for the perception and differentiation of the four facial expression categories. The categorization frequencies of expressions morphed between two prototypical expressions closely followed the choice frequencies to be expected according to the intensity of each expression category in the respective video. Moreover, increasing threat intensity was paralleled by increasing pupil size. This suggests that expressions were perceived along a continuum beyond simple template matching. This graded sensitivity may reflect the macaques’ ability to extract multifaceted information from natural facial displays, as behavioral contexts require assessment of expression strength, e.g. distinguishing mild threat from severe aggression, or evaluation of blended signals, e.g. appeasement displays despite aggressive impulses or threat displays despite fearful disposition. Random forest classification using the automatically quantified composite FACS predictors showed that the degree of jaw opening and teeth baring expectedly predicted the choice probability of the Threat and SBT categories, respectively, and revealed that lowered brows in ambiguous morphed expressions often informed the choice of the Lipsmack instead of the Neutral target, and the choice of the SBT instead of the Threat target, whereas erected ears served as a predictor for the choice of the Threat target. Under the assumption that the SBT category in our study was predominantly interpreted as a submissive signal to avoid conflict (cSBTs) (Beisner & McCowan, 2014; Flack & de Waal, 2007), lowering the brows and thus protecting the eye region against an imminent threat would be an adaptive behavior (Andrew, 1963). Conversely, raising of the brows during SBTs given as signs of formal subordination (pSBTs) could serve to enlarge the visual field for attentive monitoring of more distant possible threats, as has been implicated for human wide-eyed fear expressions (Susskind et al., 2008). However, given the restricted nature of our stimulus set and the small sample size of two monkeys, these conclusions should be taken with caution and used as the basis for further research explicitly probing and manipulating the role of eyebrow and ear movements in facial displays, which are currently underexplored in the literature. Anecdotal accounts of eyebrow movements in macaques describe lowered eyebrows combined with a staring gaze, as well as “eyebrow-flashes” (brief raising of the eyebrows) as low-grade threats, unless combined with lip-smacking (Fortman, Hewett, & Halliday, 2001; Wolfensohn & Lloyd, 2003). Rincon et al. (2023) examined the specificity of FACS AU use in rhesus macaques across different contexts and found that they raised their brows mostly in aggressive contexts and less so in submissive or affiliative situations, whereas lowered eyebrows were almost exclusively displayed in submissive contexts. Erected ears (EAU1 or EAU2) occurred in aggressive encounters, whereas flattened ears were registered in all three contexts. In a similar vein, Partan (2002) reported that pant-threat vocalizations were most frequently accompanied by forward facing ears. Humans possess little ear muscle control and thus ear movements do not play a role in facial expressions, whereas human eyebrow movements are extensive and nuanced, but do not seem diagnostic for a specific facial expression category (Mielke et al., 2022). It has been suggested that eyebrows are moved according to the momentary appropriate biological function of increasing or decreasing the visual input (Ekman, 1979, 2004; Susskind et al., 2008).

### Expression recognition transcends motion coherence, appearance degradation and species-specific form but requires knowledge of social meaning

Temporal continuity of motion was not necessary for the perception of a facial expression, as frame-scrambled prototypical avatar expressions were well recognized and elicited the same pupil reactions as temporally coherent sequences. Furthermore, fully naturalistic facial appearance was not critical, as the prototypical monkey expressions displayed by furless and colorless monkey avatars, as well as a human avatar, were still largely correctly categorized and accompanied by differential pupil sizes. The furless and colorless avatars were previously shown to be perceived as “uncanny”, causing gaze aversion and distressed facial reactions (Siebert et al., 2020). The current findings, demonstrating preserved categorization and pupil size modulation, underline that these avatars were realistic enough to be perceived as biologically plausible agents, which tend to evoke feelings of eeriness towards realistic, yet not fully realistic artificial agents (“uncanny valley”, Mori, 1970/2012). This becomes clearest in contrast to the even more unrealistic bare wireframe head avatar that did not elicit aversion in Siebert et al. (2020) and for which categorization failed in the present study, mirrored by a lack of pupil size differences between expressions. We found the same results for form-from-motion point-light expressions. This indicates that monkeys require discernable facial shape and somewhat realistic texture to interpret facial movements as meaningful facial expressions. Point-light stimuli are widely used in human research (for a review see e.g. Giese and Poggio (2003)), but behavioral discrimination studies using point-light stimuli in macaques are rare and may require training (Vangeneugden, Vancleef, Jaeggli, VanGool, & Vogels, 2010).

The relative invariance of monkey facial expressions to species-specific facial shape parallels previous findings that human observers recognize expressions displayed by both human and monkey avatars (N. Taubert et al., 2021). Both findings align with the hypothesis that facial dynamics are represented independently of facial form (Bernstein, Erez, Blank, & Yovel, 2018; Bernstein & Yovel, 2015; Duchaine & Yovel, 2015) and could be achieved via norm-referenced encoding of facial expression in terms of difference to a neutral (norm) face within a particular shape domain (Stettler, Lappe, & Giese, 2026). However, human facial expressions received unsystematic categorizations and did not elicit differential pupil reactions, suggesting that they were not mapped onto the trained categories. Under the caveat that only one monkey could be tested with human expressions, this argues against a homologous interpretation of human and macaque expressions. Categorizations followed neither presumed homologous intent (smiling/happy and lip-smacking, gasping/fearful and SBT, scowling/angry and threat), nor proposed phylogenetic homology, such as human smiles and macaque SBT displays (e.g. Kanazawa (1996), but compare Preuschoft (1992), Preuschoft and van Hooff (1995) and van Hooff (1967) for discussions on phylogenetic homology in facial displays). Moreover, the categorization pattern was not indicative of low-level feature matching, as human expressions sharing elements with monkey expressions, such as tooth exposure and retracted lip corners in smiles and SBT displays, or mouth opening in gasping (fear) expressions and threats, were not categorized accordingly. Instead, FACS-based visual similarities between human expressions and monkey expressions were uncorrelated with the target choice frequencies. This failure to categorize human expressions likely reflects limited experience, as monkeys in our facility rarely see human faces without masks, let alone have the opportunity to learn behavioral consequences associated with distinct human facial expressions. Without learned associations linking these facial configurations to behavioral intentions and predicted outcomes, human expressions remained socially meaningless to the tested macaque, eliciting neither systematic categorization nor differential arousal. These results accord with Zhu et al. (2013), who found no differential fMRI activations for human facial expressions in macaques. While some studies have used human facial expressions to probe visual encoding mechanisms of subtle facial feature changes in the macaque brain (e.g. Wehrheim, Alamooti, Ramezanpour, & Kar, 2025), our findings highlight the importance of using conspecific facial expressions for connecting visual perception to social information. Thus, we conclude that facial expression perception in the present study emerged from integrating expression-characteristic visual features with social knowledge, enabling flexible, context-sensitive interpretation of communicative signals.

## DECLARATIONS

### Ethics approval

All animal procedures were approved by the veterinary administration (Regierungspräsidium Tübingen, Abteilung Tierschutz), permit number N7/18, and complied with German law (Tierschutz-Versuchstierverordnung) and the National Institutes of Health’s Guide for the Care and Use of Laboratory Animals.

### Author contributions

R.S., M.A.G., and P.T. designed research; N.T. generated the avatar stimuli; R.S. collected data, analyzed data and wrote the initial version of the manuscript; R.S., N.T., M.A.G., and P.T. revised the manuscript.

## Acknowledgements

We are grateful to Andreas Nieder, Ziad Hafed, Markus Siegel and their respective coworkers for providing access to and letting us film their animals, which served as experimental stimuli in this study.

## Data availability

All data are available on the Open Science Framework repository at https://osf.io/y3gc8/?view_only=f17d5149e32e4a94ba3d0940b6f32b3f

## SUPPLEMENTARY

### Methods: Generation of avatars

The creation of the dynamic monkey head avatar is described in detail elsewhere (Siebert et al., 2020; N. Taubert et al., 2021). In short, the monkey head model was based on MRI scans transformed into a mesh surface, which was controlled by embedded ribbons modeled after the macaque’s facial muscles (Parr et al., 2010). The ribbon control joints corresponded to motion-captured infrared markers, which had been attached to a monkey’s face in order to record the facial movements with a VICON FX20 Motion Capture System. Human facial movements were recorded analogously from a 40-year-old male human subject. The segmented movement trajectories were preprocessed with VICON Nexus 1.85 software and time-normalized and smoothed with a generative Bayesian method combining Gaussian process latent variable models and Gaussian process dynamic models, which allows for morphing between expressions (see N. Taubert et al. (2021) Appendix 1 for details on the algorithm). The textures and skin material of the monkey avatar were hand-painted using photographic references, and additional texture maps were created in Adobe Photoshop to represent various skin layers. The fur was simulated with Autodesk Maya’s XGen Interactive Grooming tool. All movie frames were generated with Autodesk Arnold Renderer. The human avatar was based on a female face scan and texture maps provided by EISKO, with custom adjustments to the color and texture maps to match the monkey head model. The movement deformations were achieved via control points equivalent to the monkey motion-captured markers, following the human muscle anatomy (N. Taubert et al., 2021).

This resulted in a monkey head avatar able to display the prototypical, naturalistic neutral, lip-smacking, SBT and threat expressions, as well as expressions morphed across two categories in steps of 25 % intensity between each pair of prototypical expressions. Additionally, we tested scrambled-frame versions of the prototypical expressions (lacking temporal continuity of the movement), dynamic point-light expressions, which contained the same movement patterns as the prototypical naturalistic avatars, but no texture (instead white dots of 200 ms limited-lifetime distributed over the avatar surface mesh on a black background), and less realistic render types of the monkey avatar, which had been used in a previous study to test for an “uncanny valley” of aversion towards artificial agents (Siebert et al., 2020): an unnatural wireframe head without texture; a textured, but grayscale avatar; and a colored, but still furless avatar. Furthermore, we tested the monkeys’ categorization of the human avatar displaying monkey expressions, the human avatar presenting human expressions (smiling/happy, gasping/fearful, scowling/angry) and the monkey avatar showing those human expressions.

## Supplementary Tables and Figures

**Table S1.**
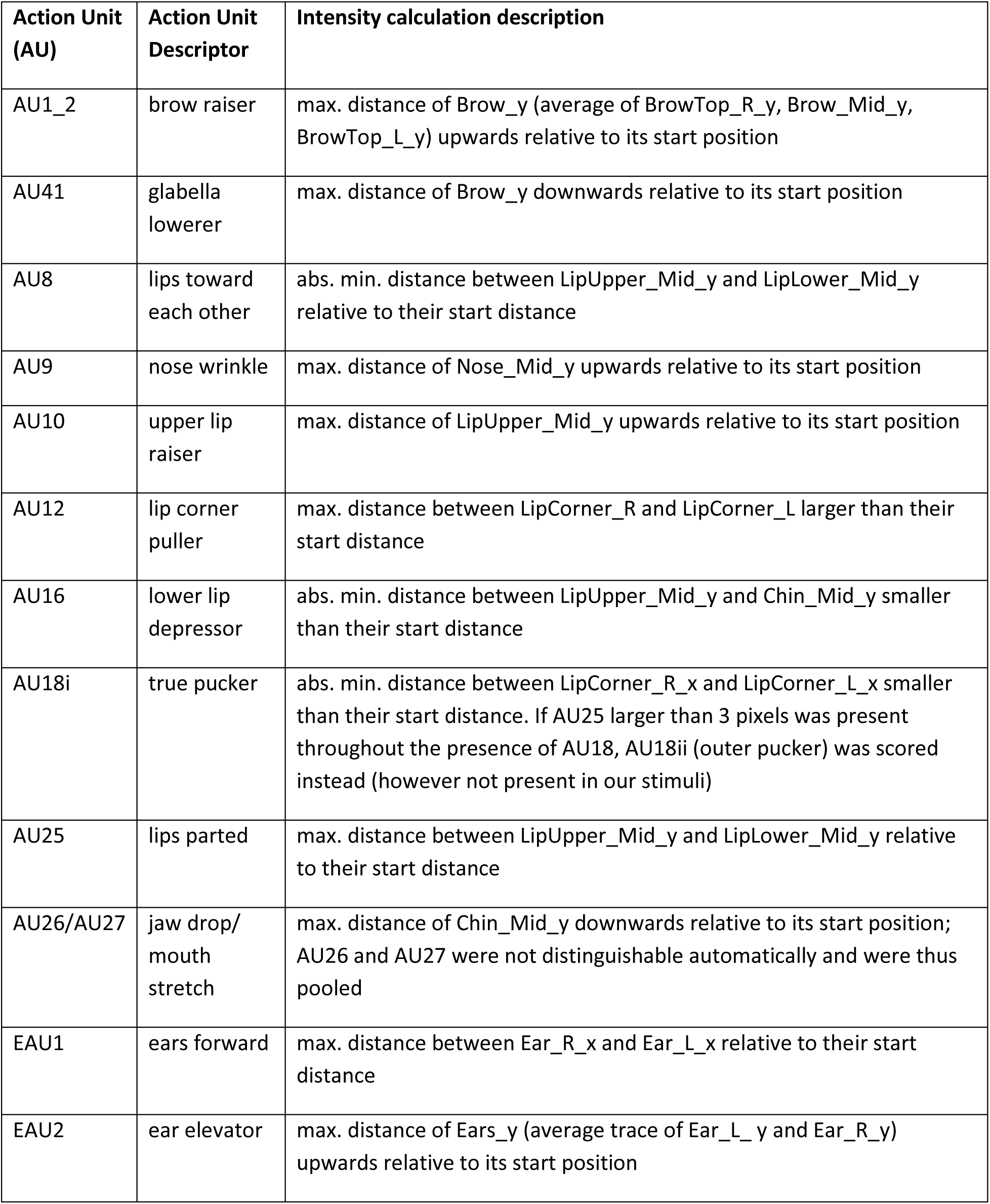
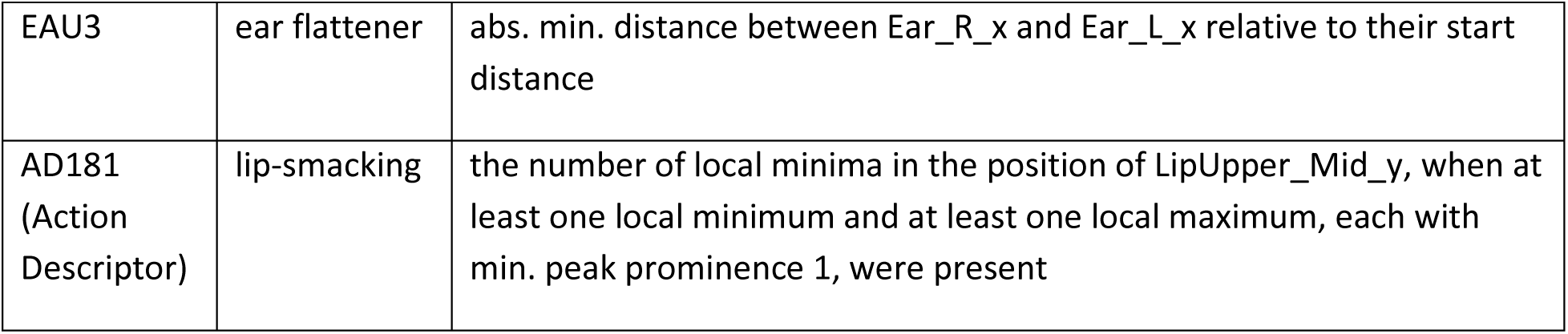
Overview of automatic MaqFACS (Macaque Facial Action Coding System) AU (Action Unit) activation and intensity calculation. Note that AU6 was not quantified, because it was not present in our stimuli and because appropriate markers are not available in the DLC primate face model

| Action Unit (AU) | Action Unit Descriptor | Intensity calculation description |
| --- | --- | --- |
| AU1_2 | brow raiser | max. distance of Brow_y (average of BrowTop_R_y, Brow_Mid_y, BrowTop_L_y) upwards relative to its start position |
| AU41 | glabella lowerer | max. distance of Brow_y downwards relative to its start position |
| AU8 | lips toward each other | abs. min. distance between LipUpper_Mid_y and LipLower_Mid_y relative to their start distance |
| AU9 | nose wrinkle | max. distance of Nose_Mid_y upwards relative to its start position |
| AU10 | upper lip raiser | max. distance of LipUpper_Mid_y upwards relative to its start position |
| AU12 | lip corner puller | max. distance between LipCorner_R and LipCorner_L larger than their start distance |
| AU16 | lower lip depressor | abs. min. distance between LipUpper_Mid_y and Chin_Mid_y smaller than their start distance |
| AU18i | true pucker | abs. min. distance between LipCorner_R_x and LipCorner_L_x smaller than their start distance. If AU25 larger than 3 pixels was present throughout the presence of AU18, AU18ii (outer pucker) was scored instead (however not present in our stimuli) |
| AU25 | lips parted | max. distance between LipUpper_Mid_y and LipLower_Mid_y relative to their start distance |
| AU26/AU27 | jaw drop/<br>mouth stretch | max. distance of Chin_Mid_y downwards relative to its start position; AU26 and AU27 were not distinguishable automatically and were thus pooled |
| EAU1 | ears forward | max. distance between Ear_R_x and Ear_L_x relative to their start distance |
| EAU2 | ear elevator | max. distance of Ears_y (average trace of Ear_L_y and Ear_R_y) upwards relative to its start position |
| EAU3 | ear flattener | abs. min. distance between Ear_R_x and Ear_L_x relative to their start distance |
| AD181<br>(Action<br>Descriptor) | lip-smacking | the number of local minima in the position of LipUpper_Mid_y, when at least one local minimum and at least one local maximum, each with min. peak prominence 1, were present |

**Fig. S1.**
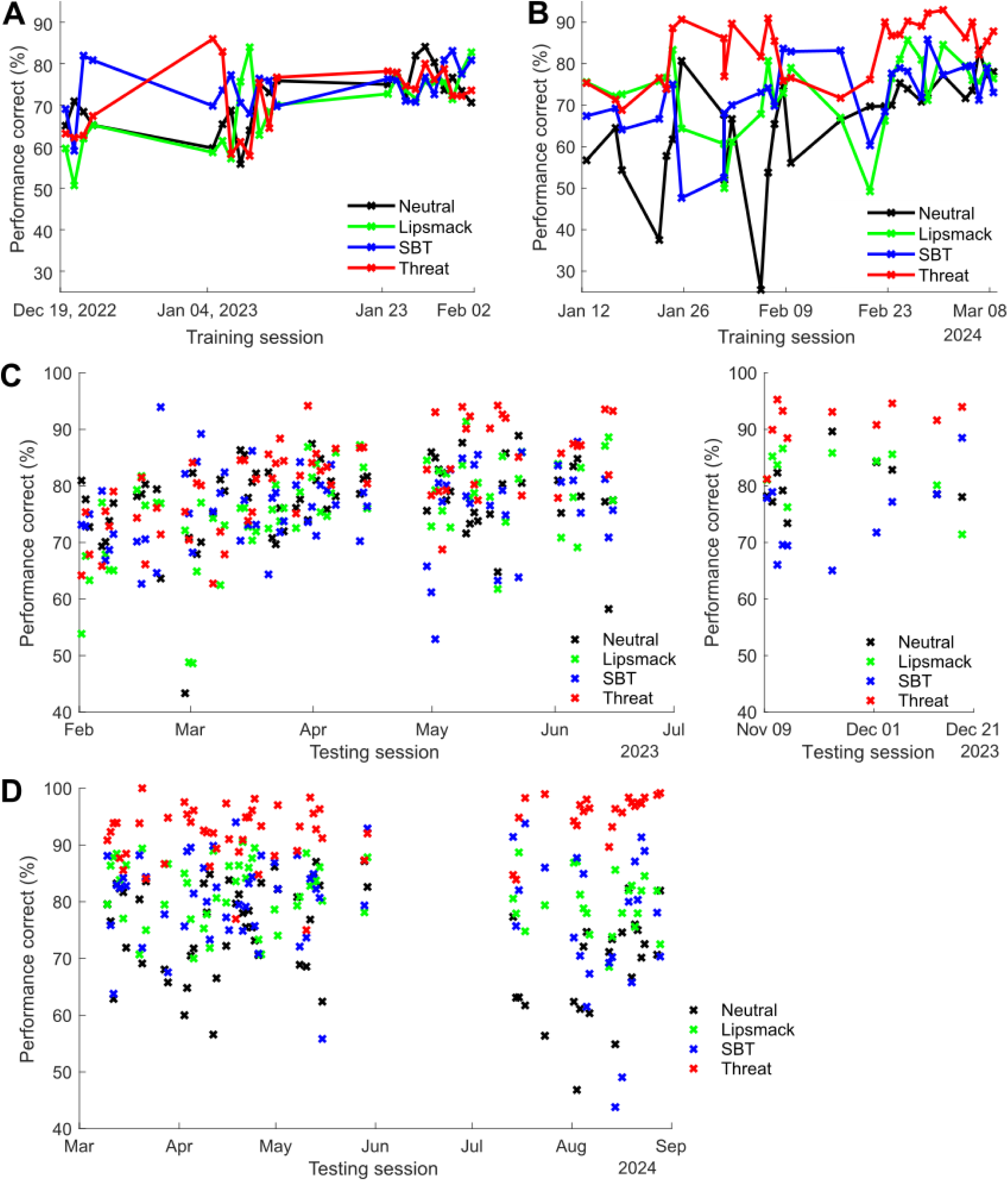
Categorization performances for the training stimuli. **(A)** Performance of monkey 1 for each expression category during the final training phase, in which trials were presented fully randomized. **(B)** Performance of monkey 2 during the final training phase. **(C)** Performance of monkey 1 on the trained videos for each test session. The left panel shows the main test phase, the right panel shows the later test phase, during which only the human expressions were tested. **(D)** Performance of monkey 2 on the trained videos for each test session.

**Fig. S2.**
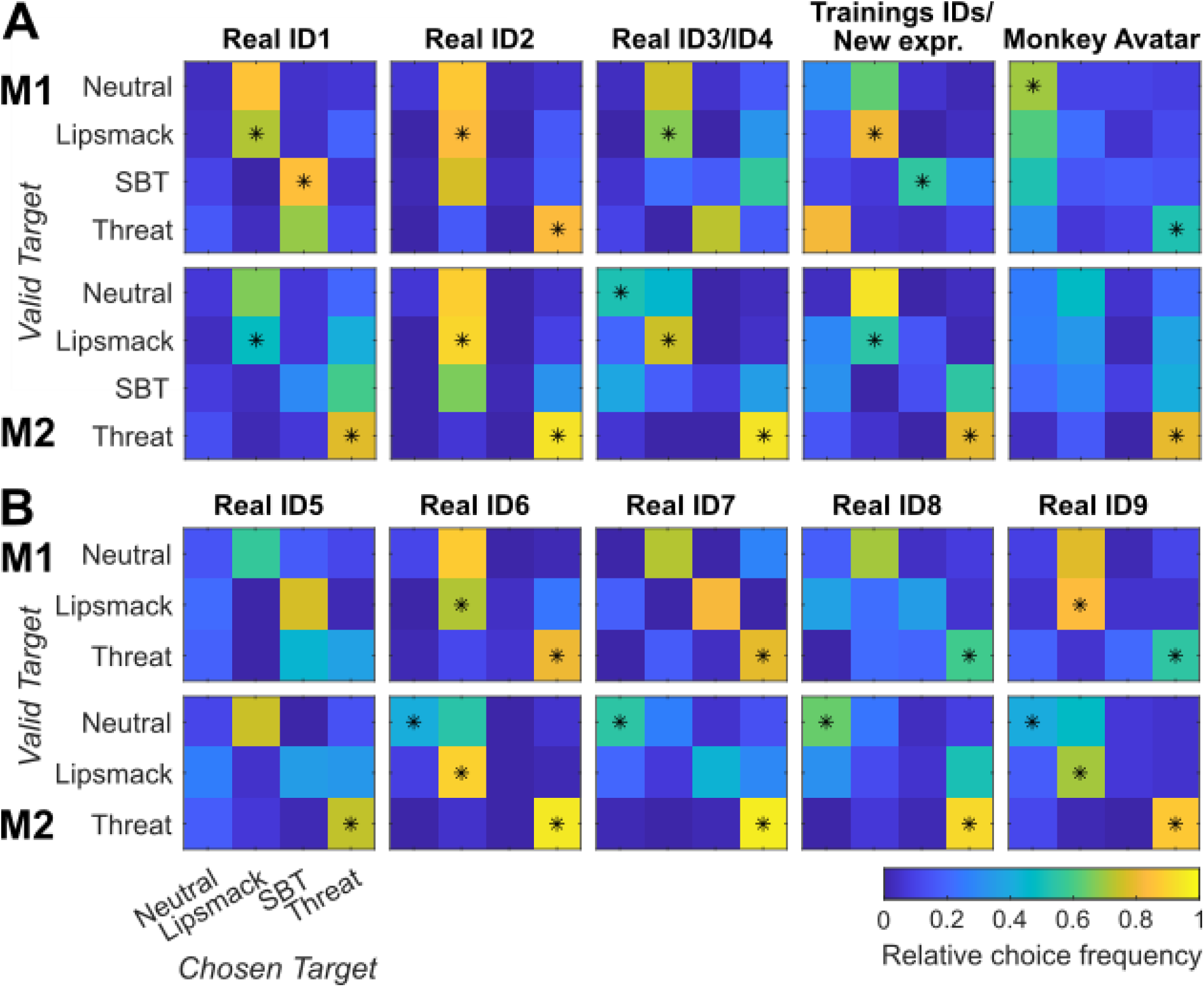
Expression categorization frequencies of all pure test expressions for monkey 1 (M1) and monkey 2 (M2), *N* per video between 50 and 60 for each stimulus identity and each subject. Asterisks indicate significant generalization (binomial tests, p < 0.05, FDR-corrected for all real monkey videos/Bonferroni-corrected for the monkey avatars). **(A)** New monkey identities (ID1-4), previously seen monkey identities with previously unseen expression categories (Training IDs/New expr.) and naturalistic monkey avatar, complete expression repertoire. **(B)** New monkey identities with incomplete expression repertoire, SBT missing (ID 5-9)

**Fig. S3.**
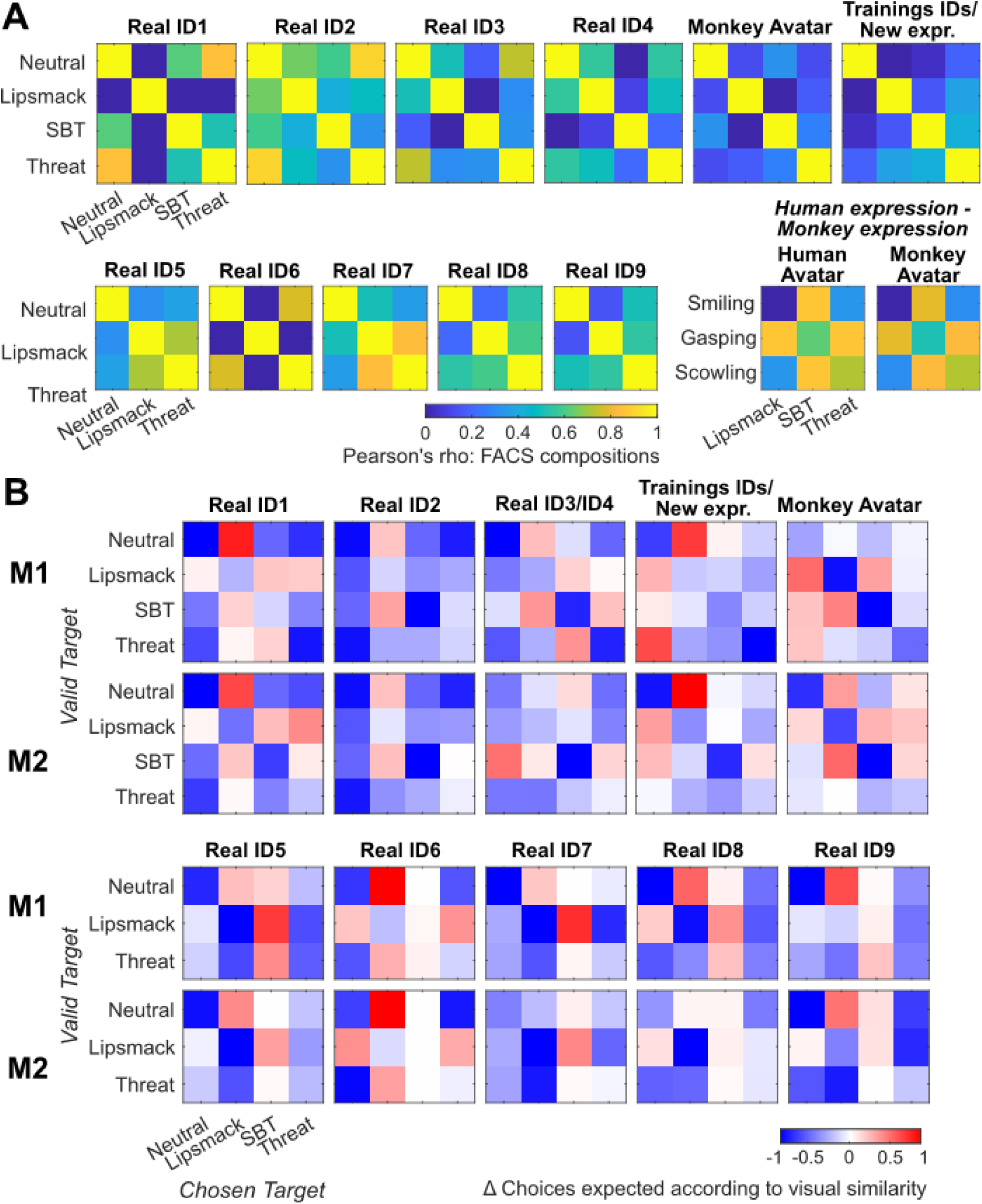
**(A)** Visual similarity matrices per test ID for all pure test expressions and visual similarity matrices between human expressions and monkey expressions. Visual similarity was calculated as the Pearson correlation between the quantified FACS compositions of an expression pair. **(B)** Difference matrices per test ID for all pure test expressions, for monkey 1 (M1) and monkey 2 (M2) separately. Differences were calculated between observed relative choice frequency and the expected choice frequency according to the visual similarity between the test expression and the expression associated with the selected target. A value of zero (white) indicates that the target was selected as often as predicted by the visual similarity, positive values (red) indicate that the target was selected more often, and negative values (blue) indicate that the target was selected less often

**Fig. S4.**
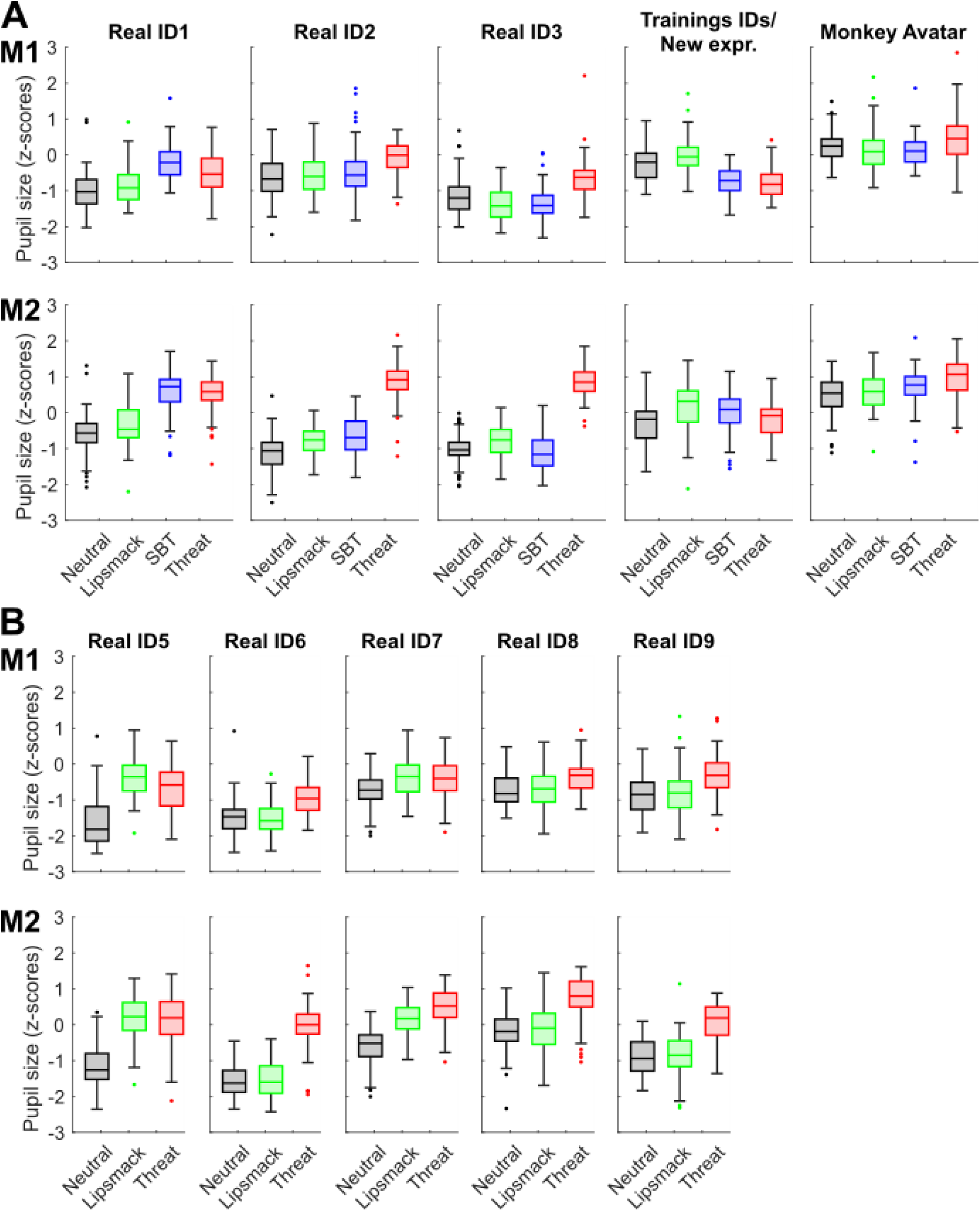
Boxplots of pupil sizes during the viewing of each expression category for monkey 1 (M1) and monkey 2 (M2). **(A)** N New monkey identities (ID1-4), previously seen monkey identities with previously unseen expression categories (Training IDs/New expr.) and naturalistic monkey avatar, complete expression repertoire **(B)** New monkey identities with incomplete expression repertoire, SBT missing (ID 5-9)

**Table S2.** Multinomial Logistic Regression (MLR) model comparisons for categorization choices of the pure (real and naturalistic monkey avatar) expression videos (*N* = 4086)

| MLR Model Nr. | Predictors [number of predictors] | Adj. $R^2$ (explained variance) | Deviance $D$ (from a perfect model) | $\Delta$ Deviance $D$, $p$ | $\chi^2$, df, $p$ |
| --- | --- | --- | --- | --- | --- |
| 1 | Expression [1] | 0.1476 | 9039.40 | – | 1593.55, 9, $p < 0.0001$ |
| 2 | Pupil size [1] | 0.0582 | 10001.65 | – | 631.30, 3, $p < 0.0001$ |
| 3 | Expression + Identity [2] | 0.2517 | 7854.96 | 1184.44, $p < 0.0001$ ($\Delta$ to Model 1) | 2777.99, 48, $p < 0.0001$ |
| 4 | FACS (LipPucker, TeethBare, JawOpen, BrowRaise, BrowLow, EarsErect, EarsFlat) [7] | 0.1445 | 9048.91 | – | 1584.04, 21, $p < 0.0001$ |
| 5 | FACSx: FACS + 3-way interactions between mouth predictors (LipPucker, TeethBare, JawOpen) x brow predictors (BrowRaise, BrowLow) x ear predictors (EarsErect, EarsFlat) [19] | 0.2262 | 8108.27 | 940.64, $p < 0.0001$ ($\Delta$ to Model 4) | 2524.68, 57, $p < 0.0001$ |
| 6 | FACSx + Weight [20] | 0.4430 | 5796.31 | 2311.96, $p < 0.0001$ ($\Delta$ to Model 5) | 4836.64, 60, $p < 0.0001$ |
| 7 | FACSx + Weight + Avert [21] | 0.4859 | 5334.73 | 461.58, $p < 0.0001$ ($\Delta$ to Model 6) | 5298.22, 63, $p < 0.0001$ |
| 8 | FACSx + Weight + Avert + Subject [22] | 0.5087 | 5086.32 | 248.41, $p < 0.0001$ ($\Delta$ to Model 7) | 5546.63, 66, $p < 0.0001$ |

**Table S3.** MLR Model 8 for the categorizations of the pure (real and naturalistic monkey avatar) expressions. Coefficient names, log odds values, standard errors and t-statistics with p-values.

| Coefficient | Value | Standard error | t-statistic | p-value |
| --- | --- | --- | --- | --- |
| (Intercept_TargetLipsmack) | 3.40 | 1.00 | 3.38 | 0.00072 |
| Subject_M2_TargetLipsmack | -0.91 | 0.14 | -6.41 | < 0.0001 |
| Weight_TargetLipsmack | 0.092 | 0.12 | 0.78 | 0.44 |
| BrowRaise_TargetLipsmack | -0.24 | 0.15 | -1.56 | 0.12 |
| BrowLow_TargetLipsmack | -1.14 | 2.63 | -0.44 | 0.66 |
| TeethBare_TargetLipsmack | -5.76 | 1.39 | -4.15 | < 0.0001 |
| LipPucker_TargetLipsmack | 2.88 | 1.11 | 2.59 | 0.0095 |
| JawOpen_TargetLipsmack | 7.36 | 1.52 | 4.84 | < 0.0001 |
| EarsErect_TargetLipsmack | -3.65 | 0.97 | -3.77 | 0.00016 |
| EarsFlat_TargetLipsmack | 5.99 | 2.25 | 2.67 | 0.0076 |
| Avert_TargetLipsmack | -4.79 | 0.46 | -10.45 | < 0.0001 |
| BrowRaise:TeethBare:EarsErect_TargetLipsmack | -8.25 | 1.89 | -4.36 | < 0.0001 |
| BrowLow:TeethBare:EarsErect_TargetLipsmack | -11.99 | 3.77 | 3.18 | 0.0015 |
| BrowRaise:LipPucker:EarsErect_TargetLipsmack | 15.88 | 3.42 | 4.95 | < 0.0001 |
| BrowLow:LipPucker:EarsErect_TargetLipsmack | -21.26 | 7.76 | -2.74 | 0.0062 |
| BrowRaise:JawOpen:EarsErect_TargetLipsmack | 8.88 | 2.32 | 3.83 | 0.00013 |
| BrowLow:JawOpen:EarsErect_TargetLipsmack | -19.32 | 7.58 | -2.55 | 0.011 |
| BrowRaise:TeethBare:EarsFlat_TargetLipsmack | 9.67 | 2.32 | 4.16 | < 0.0001 |
| BrowLow:TeethBare:EarsFlat_TargetLipsmack | 40.17 | 13.33 | 3.01 | 0.0026 |
| BrowRaise:LipPucker:EarsFlat_TargetLipsmack | -14.41 | 3.41 | -4.23 | < 0.0001 |
| BrowLow:LipPucker:EarsFlat_TargetLipsmack | 15.13 | 4.63 | 3.27 | 0.0011 |
| BrowRaise:JawOpen:EarsFlat_TargetLipsmack | -12.53 | 2.73 | -4.58 | < 0.0001 |
| BrowLow:JawOpen:EarsFlat_TargetLipsmack | -45.57 | 11.37 | -4.01 | < 0.0001 |
| (Intercept_TargetSBT) | 10.55 | 2.07 | 5.09 | < 0.0001 |
| Subject_M2_TargetSBT | -2.07 | 0.19 | -10.81 | < 0.0001 |
| Weight_TargetSBT | -2.05 | 0.37 | -5.59 | < 0.0001 |
| BrowRaise_TargetSBT | -5.91 | 0.98 | -6.03 | < 0.0001 |
| BrowLow_TargetSBT | 5.46 | 3.95 | 1.38 | 0.17 |
| TeethBare_TargetSBT | -12.69 | 1.94 | -6.56 | < 0.0001 |
| LipPucker_TargetSBT | 0.95 | 1.32 | 0.72 | 0.47 |
| JawOpen_TargetSBT | 14.16 | 2.00 | 7.08 | < 0.0001 |
| EarsErect_TargetSBT | 9.45 | 1.81 | 5.22 | < 0.0001 |
| EarsFlat_TargetSBT | 24.31 | 4.20 | 5.79 | < 0.0001 |
| Avert_TargetSBT | 3.05 | 0.81 | 3.77 | < 0.0001 |
| BrowRaise:TeethBare:EarsErect_TargetSBT | -8.82 | 1.30 | -6.77 | < 0.0001 |
| BrowLow:TeethBare:EarsErect_TargetSBT | -2.82 | 2.89 | -0.98 | 0.33 |
| BrowRaise:LipPucker:EarsErect_TargetSBT | 52.33 | 6.91 | 7.57 | < 0.0001 |
| BrowLow:LipPucker:EarsErect_TargetSBT | -30.80 | 10.71 | -2.88 | 0.0040 |
| BrowRaise:JawOpen:EarsErect_TargetSBT | 36.23 | 4.63 | 7.82 | < 0.0001 |
| BrowLow:JawOpen:EarsErect_TargetSBT | 11.65 | 8.30 | 1.40 | 0.16 |
| BrowRaise:TeethBare:EarsFlat_TargetSBT | 7.46 | 2.21 | 3.38 | < 0.0001 |
| BrowLow:TeethBare:EarsFlat_TargetSBT | 108.11 | 16.29 | 6.64 | < 0.0001 |
| BrowRaise:LipPucker:EarsFlat_TargetSBT | -23.11 | 5.74 | -4.03 | < 0.0001 |
| BrowLow:LipPucker:EarsFlat_TargetSBT | -8.39 | 7.46 | -1.12 | 0.26 |
| BrowRaise:JawOpen:EarsFlat_TargetSBT | -35.82 | 4.84 | -7.41 | < 0.0001 |
| BrowLow:JawOpen:EarsFlat_TargetSBT | -128.50 | 18.07 | -7.11 | < 0.0001 |
| (Intercept_TargetThreat) | 0.68 | 0.91 | 0.74 | 0.46 |
| Subject_M2_TargetThreat | 0.083 | 0.15 | 0.55 | 0.58 |
| Weight_TargetThreat | -0.52 | 0.16 | -3.26 | 0.0011 |
| BrowRaise_TargetThreat | -1.22 | 0.22 | -5.67 | < 0.0001 |
| BrowLow_TargetThreat | -11.71 | 2.49 | -4.71 | < 0.0001 |
| TeethBare_TargetThreat | -9.26 | 1.37 | -6.77 | < 0.0001 |
| LipPucker_TargetThreat | 0.50 | 0.99 | 0.51 | 0.61 |
| JawOpen_TargetThreat | 10.26 | 1.51 | 6.81 | < 0.0001 |
| EarsErect_TargetThreat | -0.0014 | 0.72 | -0.0019 | 0.99 |
| EarsFlat_TargetThreat | 6.54 | 2.01 | 3.26 | 0.0011 |
| Avert_TargetThreat | -2.15 | 0.42 | -5.18 | < 0.0001 |
| BrowRaise:TeethBare:EarsErect_TargetThreat | -5.94 | 0.83 | -7.20 | < 0.0001 |
| BrowLow:TeethBare:EarsErect_TargetThreat | 3.57 | 2.30 | 1.55 | 0.12 |
| BrowRaise:LipPucker:EarsErect_TargetThreat | 23.44 | 3.55 | 6.61 | < 0.0001 |
| BrowLow:LipPucker:EarsErect_TargetThreat | -15.90 | 7.44 | -2.14 | 0.033 |
| BrowRaise:JawOpen:EarsErect_TargetThreat | 16.57 | 2.38 | 6.97 | < 0.0001 |
| BrowLow:JawOpen:EarsErect_TargetThreat | -8.16 | 6.74 | -1.21 | 0.23 |
| BrowRaise:TeethBare:EarsFlat_TargetThreat | 6.03 | 1.41 | 4.27 | < 0.0001 |
| BrowLow:TeethBare:EarsFlat_TargetThreat | 86.15 | 12.42 | 6.94 | < 0.0001 |
| BrowRaise:LipPucker:EarsFlat_TargetThreat | -20.28 | 3.51 | -5.78 | < 0.0001 |
| BrowLow:LipPucker:EarsFlat_TargetThreat | 22.74 | 4.79 | 4.75 | < 0.0001 |
| BrowRaise:JawOpen:EarsFlat_TargetThreat | -17.22 | 2.76 | -6.23 | < 0.0001 |
| BrowLow:JawOpen:EarsFlat_TargetThreat | -65.53 | 10.66 | -6.15 | < 0.0001 |

**Table S4.**
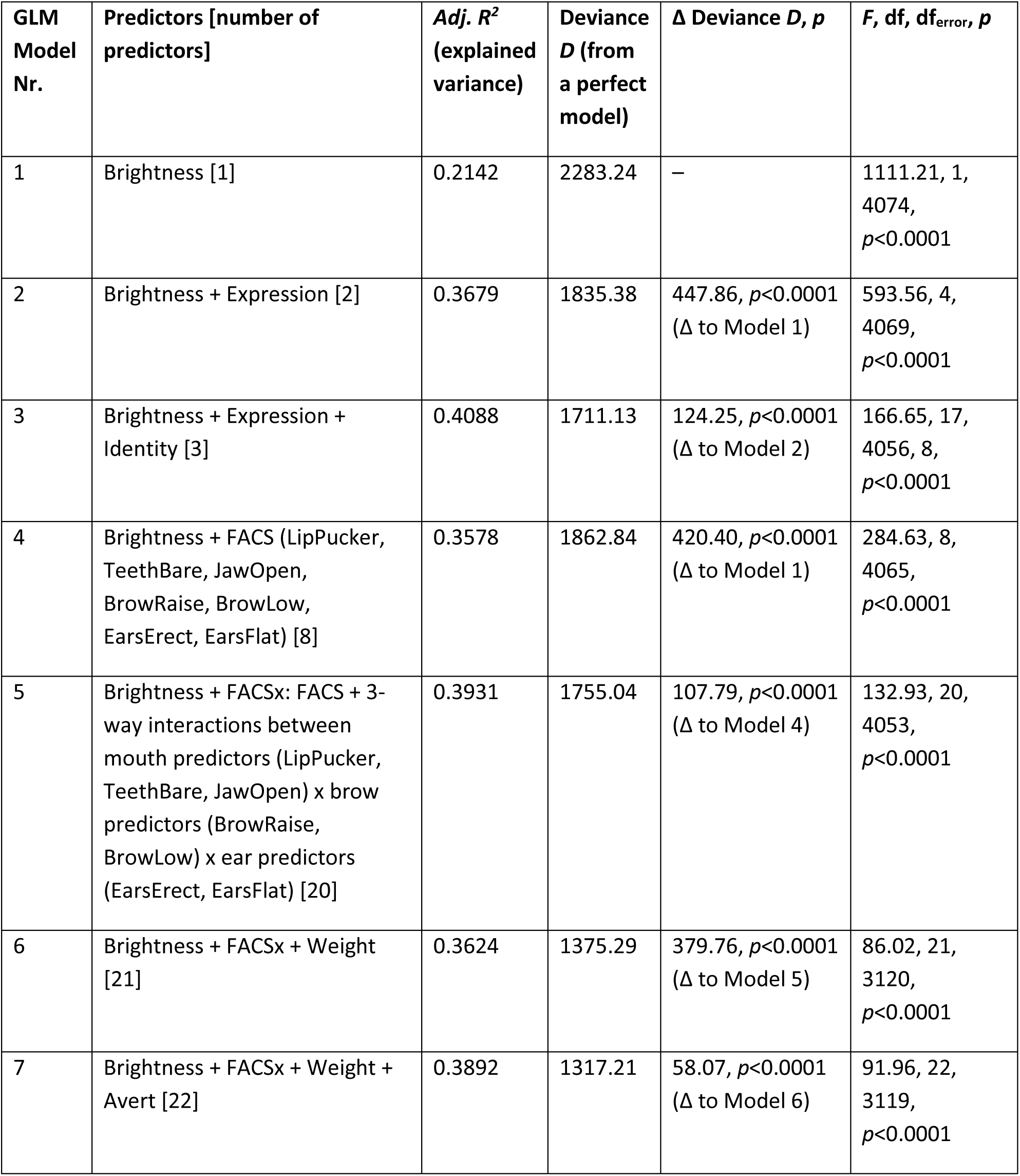
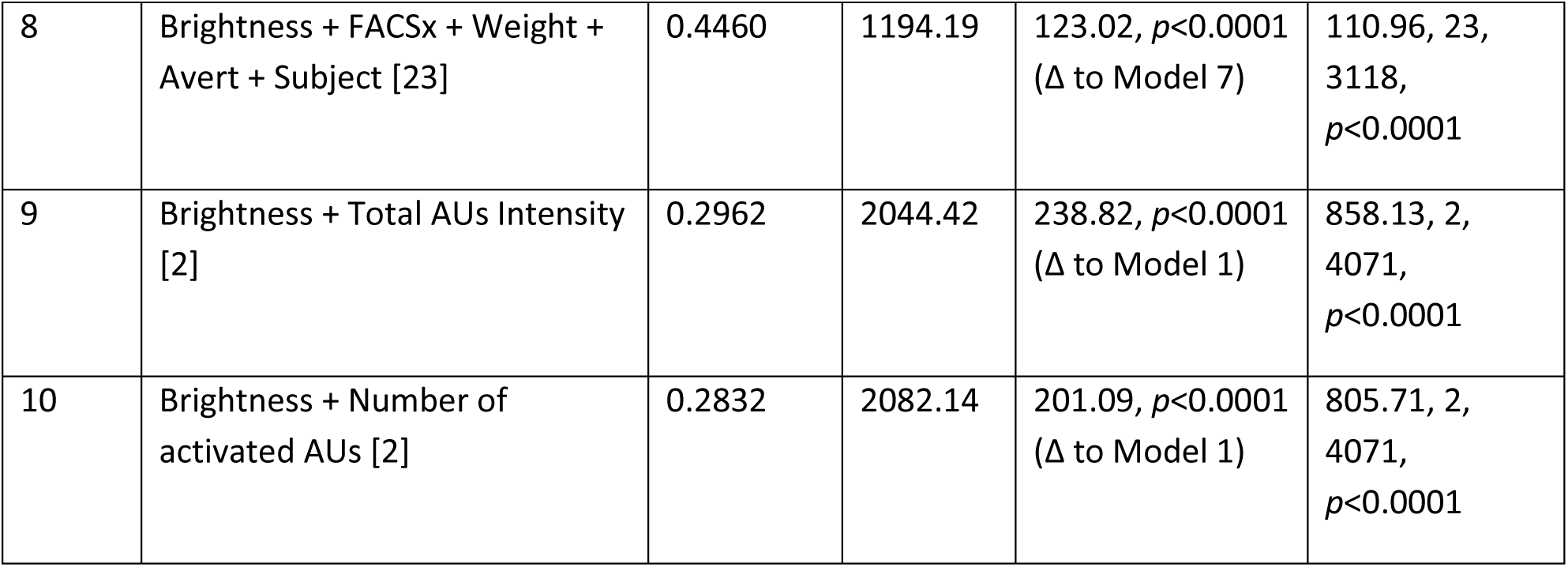
General Linear Model (GLM) model comparisons for pupil size when viewing the pure (real and naturalistic monkey avatar) expression videos (*N* = 4074)

**Fig. S5.**
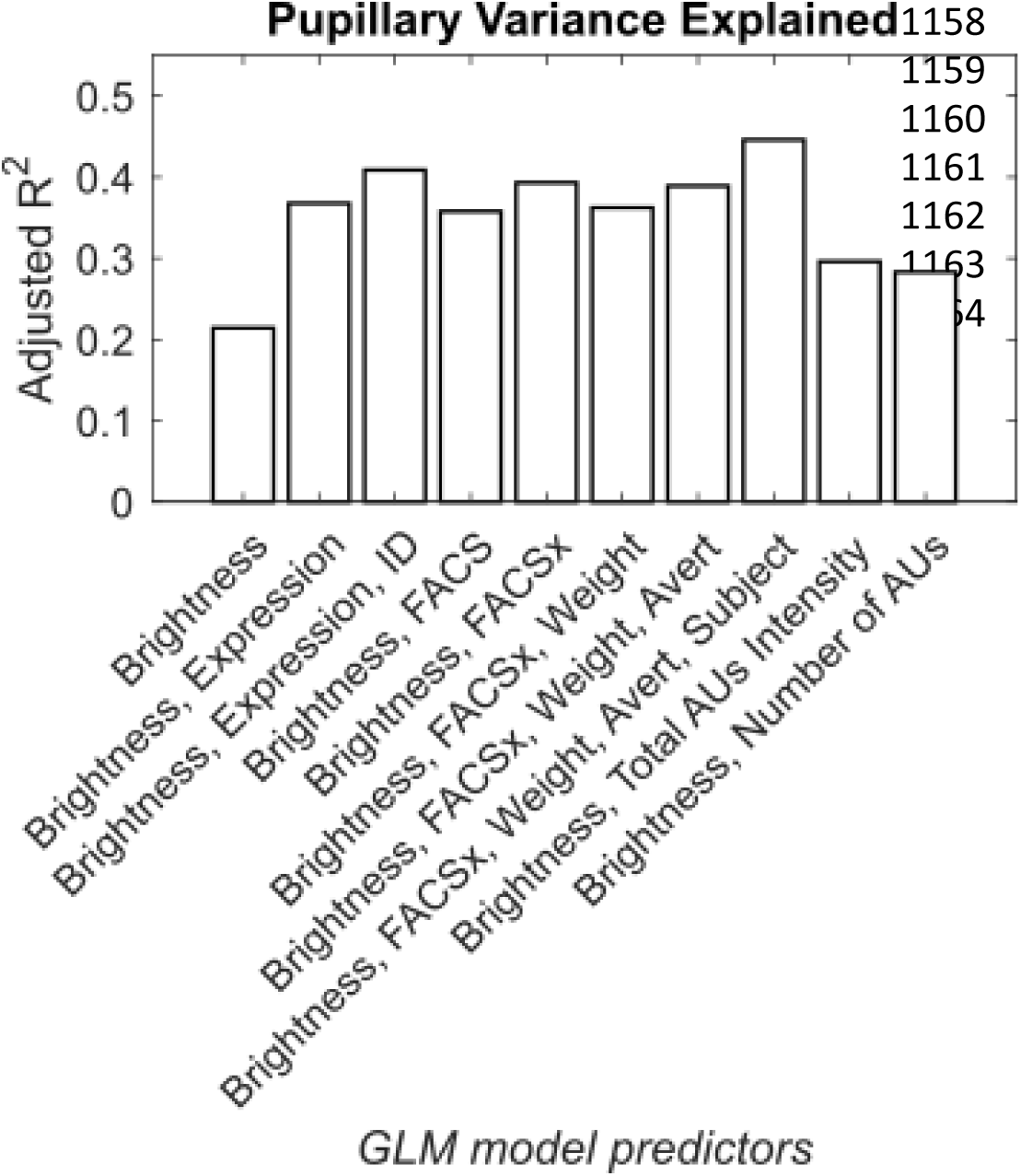
Proportion of the variance in pupil size when viewing the pure (real and naturalistic monkey avatar) expression videos explained (adjusted R^2^) by GLM models with different sets of predictors. “FACSx” stands for the FACS predictors + 3-way interactions between mouth predictors x brow predictors x ear predictors

**Fig. S6.**
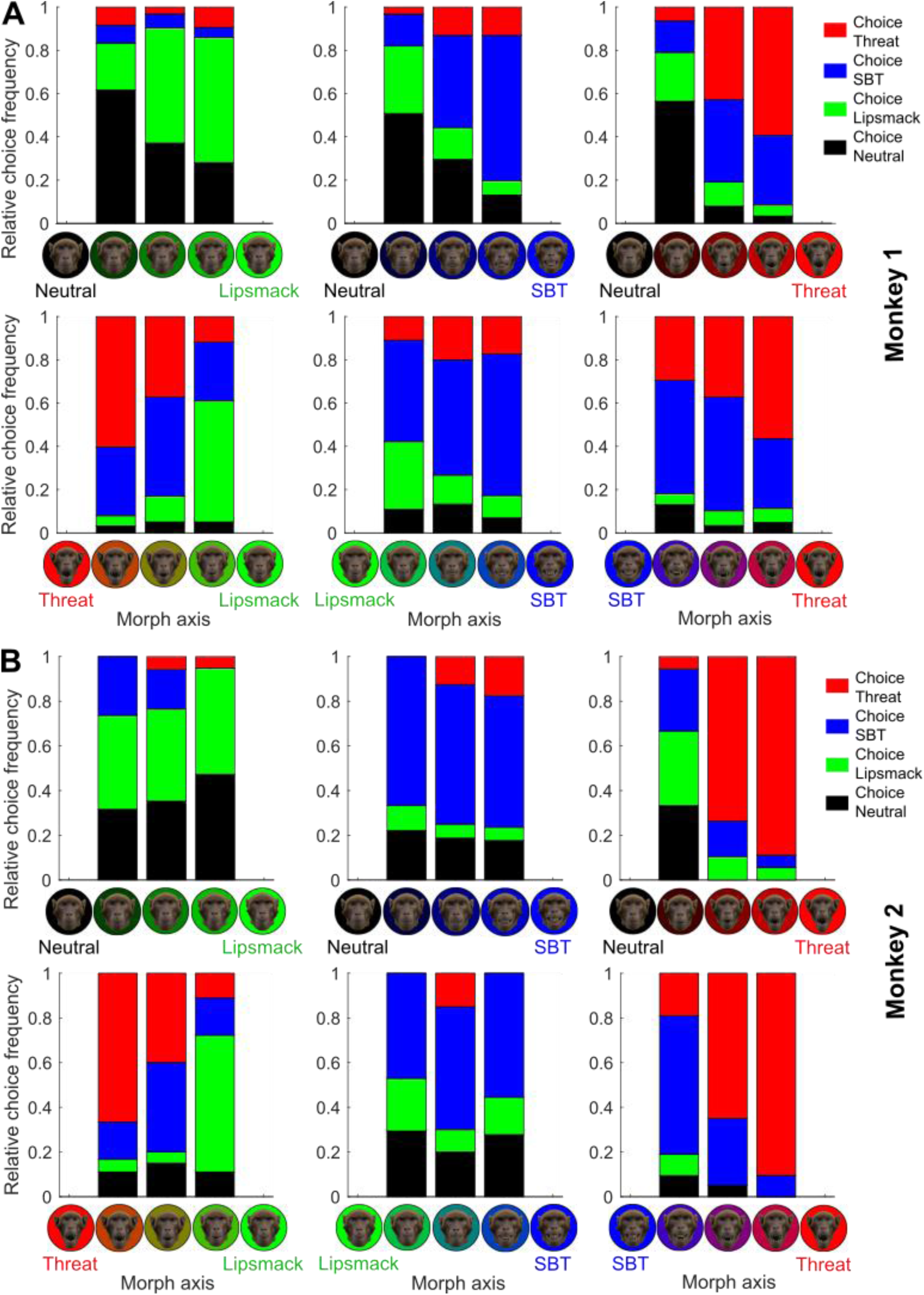
Relative choice frequencies for all morphed avatar videos, shown for each morph axis between the prototypical expressions Neutral, Lipsmack, SBT and Threat separately for monkey 1 **(A)**, *N* per video = 53, and monkey 2 **(B)**, *N* per video = 18

**Table S5.** Multinomial Logistic Regression (MLR) model comparisons for categorization choices of the morphed avatar expressions (*N* = 1432)

| Morphs<br>MLR<br>Model<br>Nr. | Predictors [number of predictors] | Adj. $R^2$<br>(explained<br>variance) | Deviance<br>$D$ (from a<br>perfect<br>model) | $\Delta$ Deviance $D$,<br>$p$ | $\chi^2$, df, $p$ |
| --- | --- | --- | --- | --- | --- |
| 1 | Lipsmack, SBT and Threat intensities [3] | 0.1756 | 3180.25 | – | 706.41, 9,<br>$p < 0.0001$ |
| 2 | Lipsmack, SBT and Threat intensities + 2-way interactions [6] | 0.1782 | 3151.92 | 28.33, $p < 0.001$<br>($\Delta$ to Model 1) | 734.73, 18,<br>$p < 0.0001$ |
| 3 | FACS (LipPucker, TeethBare, JawOpen, BrowRaise, BrowLow, EarsErect, EarsFlat) [7] | 0.1734 | 3164.55 | – | 722.11, 21,<br>$p < 0.0001$ |
| 4 | FACS + Subject [8] | 0.1729 | 3160.75 | 3.80, $p = 0.051$<br>($\Delta$ to Model 3) | 725.90, 24,<br>$p < 0.0001$ |

**Table S6.** General Linear Model (GLM) model comparisons for pupil size when viewing the morphed avatar expressions (*N* = 1431)

| Morphs<br>GLM<br>Model<br>Nr. | Predictors [number of<br>predictors] | Adj. $R^2$<br>(explained<br>variance) | Deviance<br>$D$ (from<br>a perfect<br>model) | $\Delta$ Deviance $D$, $p$ | $F$, df,<br>df<sub>error</sub>, $p$ |
| --- | --- | --- | --- | --- | --- |
| 1 | Brightness [1] | 0.0261 | 683.71 | – | 39.30, 1,<br>1429,<br>$p < 0.0001$ |
| 2 | Brightness + Lipsmack, SBT and Threat intensity [4] | 0.0876 | 639.19 | 44.52, $p < 0.0001$<br>($\Delta$ to Model 1) | 35.32, 4,<br>1426,<br>$p < 0.0001$ |
| 3 | Brightness + Lipsmack, SBT and Threat intensities + 2-way interactions [7] | 0.0884 | 637.27 | 1.92, $p = 0.59$ ($\Delta$ to Model 2) | 20.81, 7,<br>1423,<br>$p < 0.0001$ |
| 4 | Brightness + FACS (LipPucker, TeethBare, JawOpen, BrowRaise, BrowLow, EarsErect, EarsFlat) [8] | 0.0908 | 635.15 | 48.55, $p<0.0001$ ($\Delta$ to Model 1) | 18.85, 8, 1422, $p<0.0001$ |
| 5 | Brightness + FACS + Subject [9] | 0.1053 | 624.57 | 10.59, $p=0.001$ ($\Delta$ to Model 4) | 19.70, 9, 1421, $p<0.0001$ |
| 6 | Brightness + Total AUs Intensity [2] | 0.0309 | 679.87 | 3.84, $p=0.050$ ($\Delta$ to Model 1) | 23.78, 2, 1428, $p<0.0001$ |
| 7 | Brightness + Number of activated AUs [2] | 0.0320 | 679.06 | 4.65, $p=0.031$ ($\Delta$ to Model 1) | 24.66, 2, 1428, $p<0.0001$ |

**Fig. S7.**
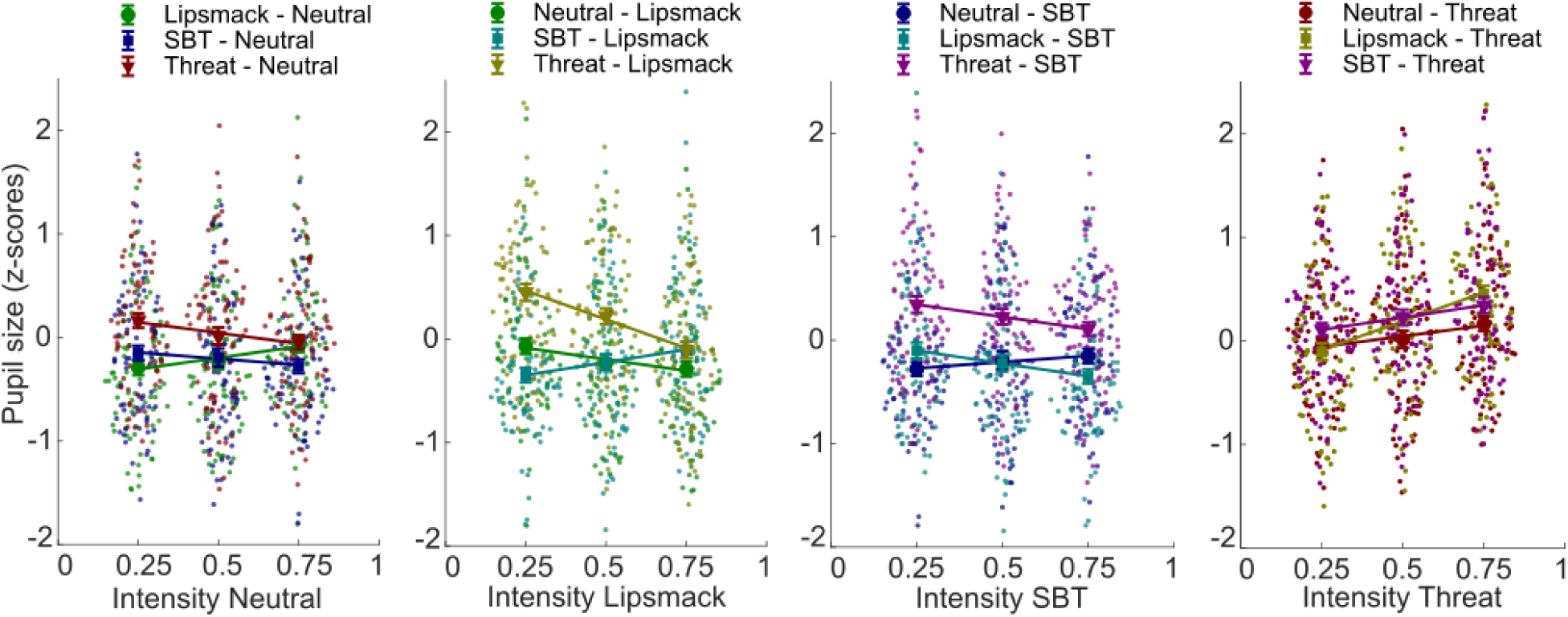
Swarm plots show pupil sizes of both monkeys during the viewing of the morphed avatar expressions, separated by expression intensity and color-coded by morph axis. Averages for each morph axis are plotted as large symbols with SEM error bars and linear trend lines are overlaid

**Fig. S8.**
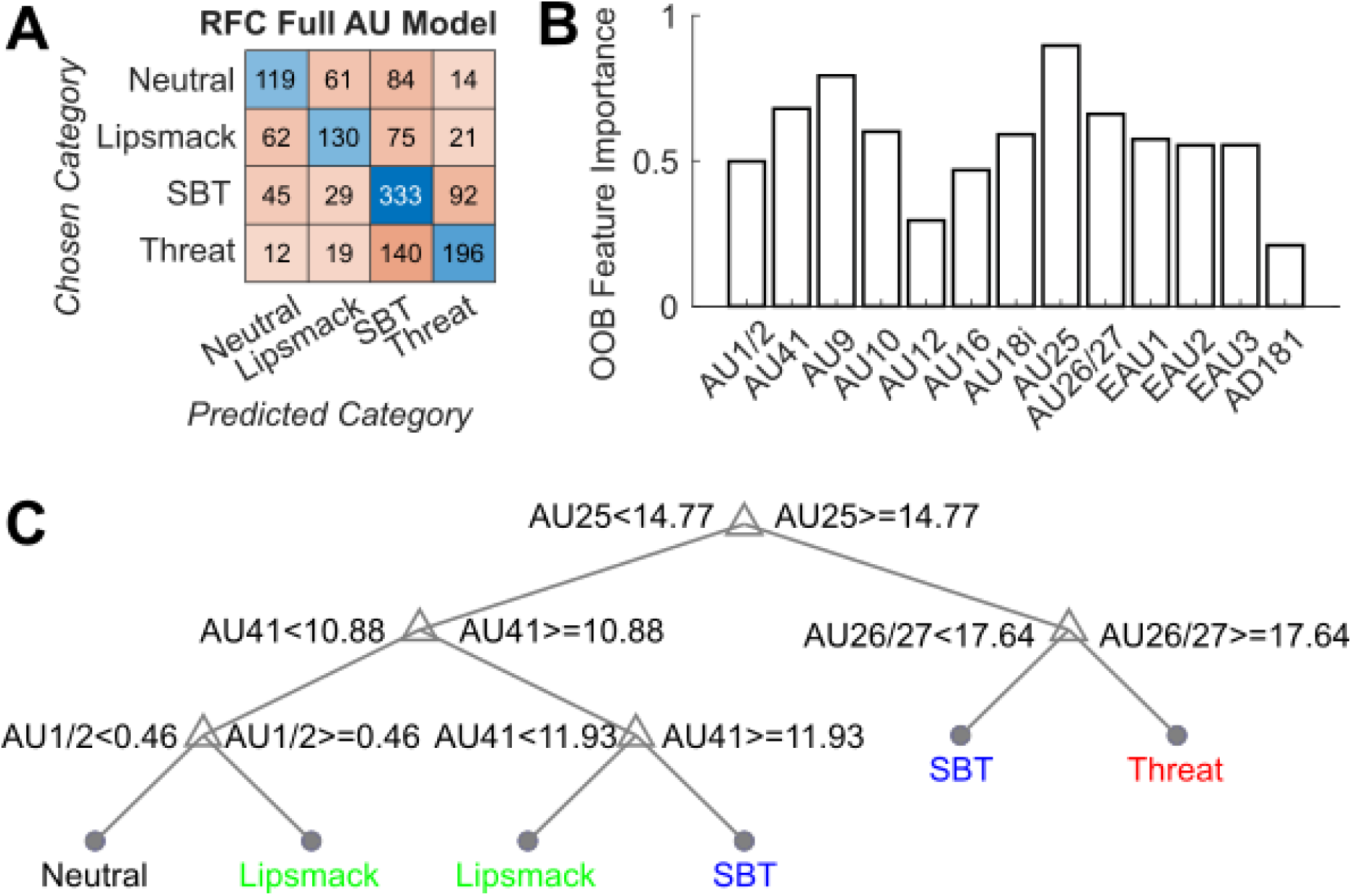
Random Forest Classification results of the morphed avatar expressions using the full set of FACS AUs as predictors (full AU model). **(A)** Heatmap showing the predicted category vs. the chosen category. **(B)** Out-of-bag feature importance of the predictors. **(C)** Simplified decision tree

**Fig. S9.**
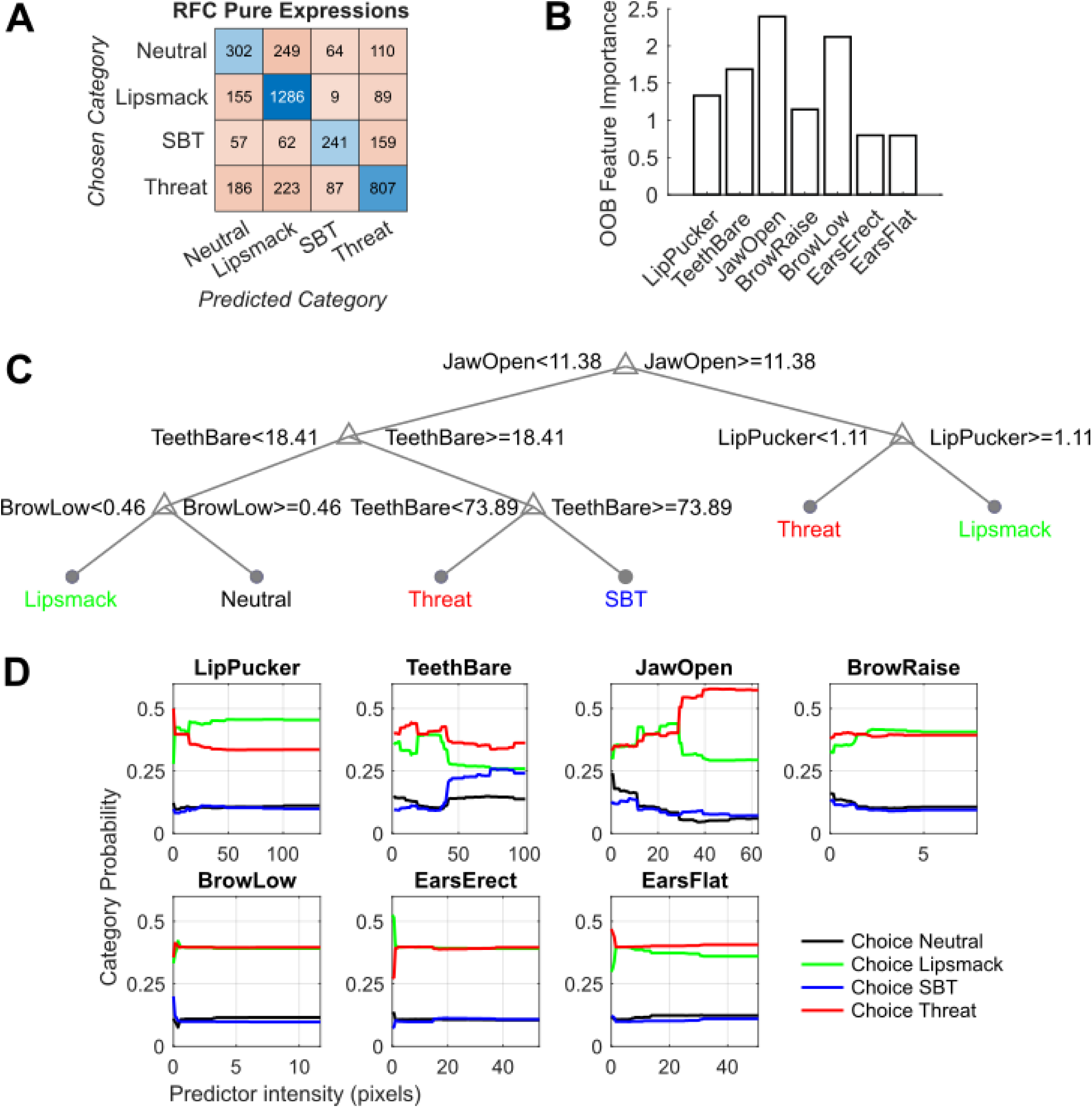
Random Forest Classification results of the pure (unambiguous) real and naturalistic avatar expressions using the composite FACS predictors (composite model). **(A)** Heatmap showing the predicted category vs. the chosen category. **(B)** Out-of-bag feature importance of the predictors. **(C)** Simplified decision tree. **(D)** Partial dependency plots

**Fig. S10.**
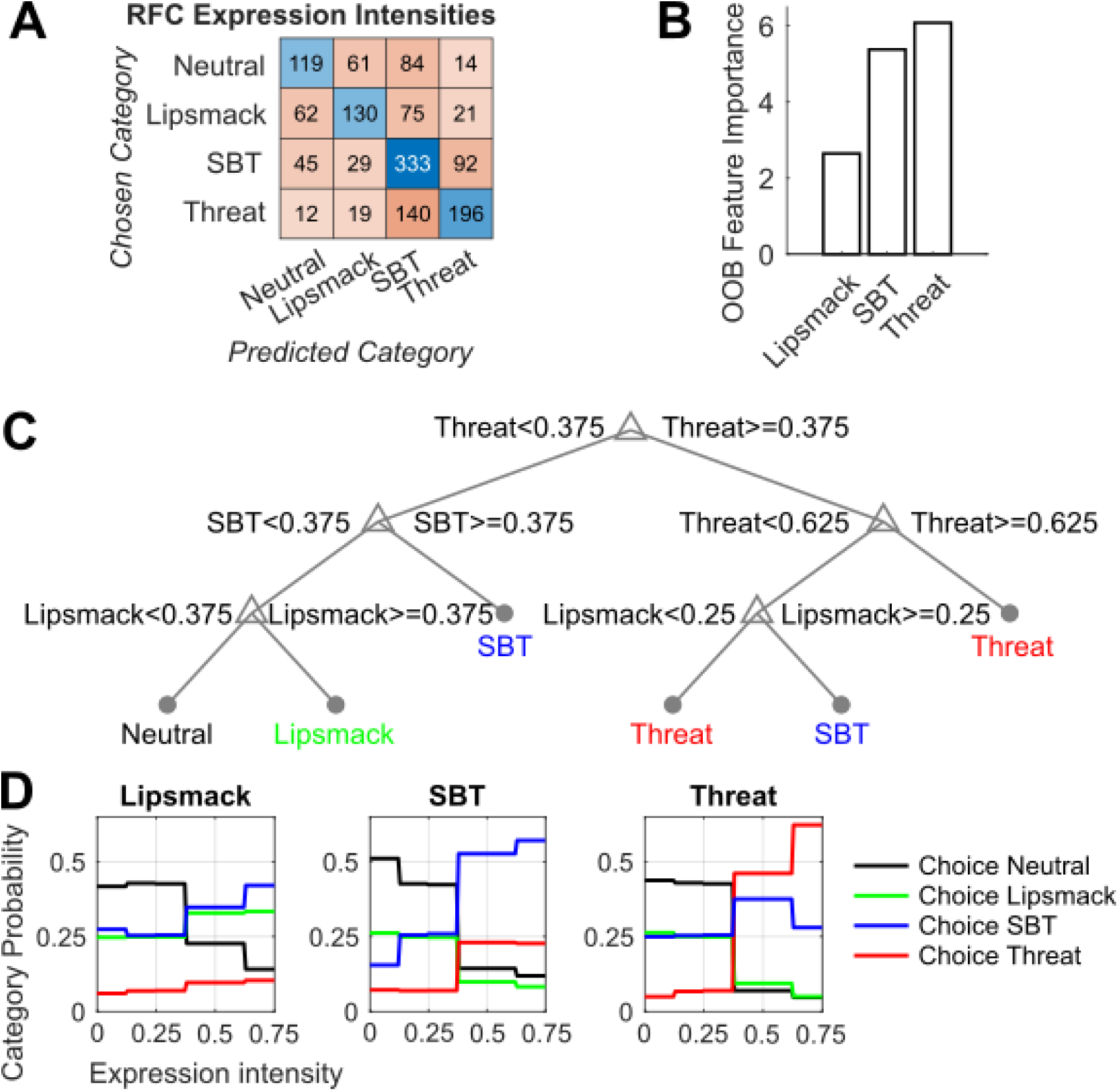
Random Forest Classification results of the morphed avatar expressions using the expression intensities in the morphs as predictors. **(A)** Heatmap showing the predicted category vs. the chosen category. **(B)** Out-of-bag feature importance of the predictors. **(C)** Simplified decision tree. **(D)** Partial dependency plots

**Fig. S11.**
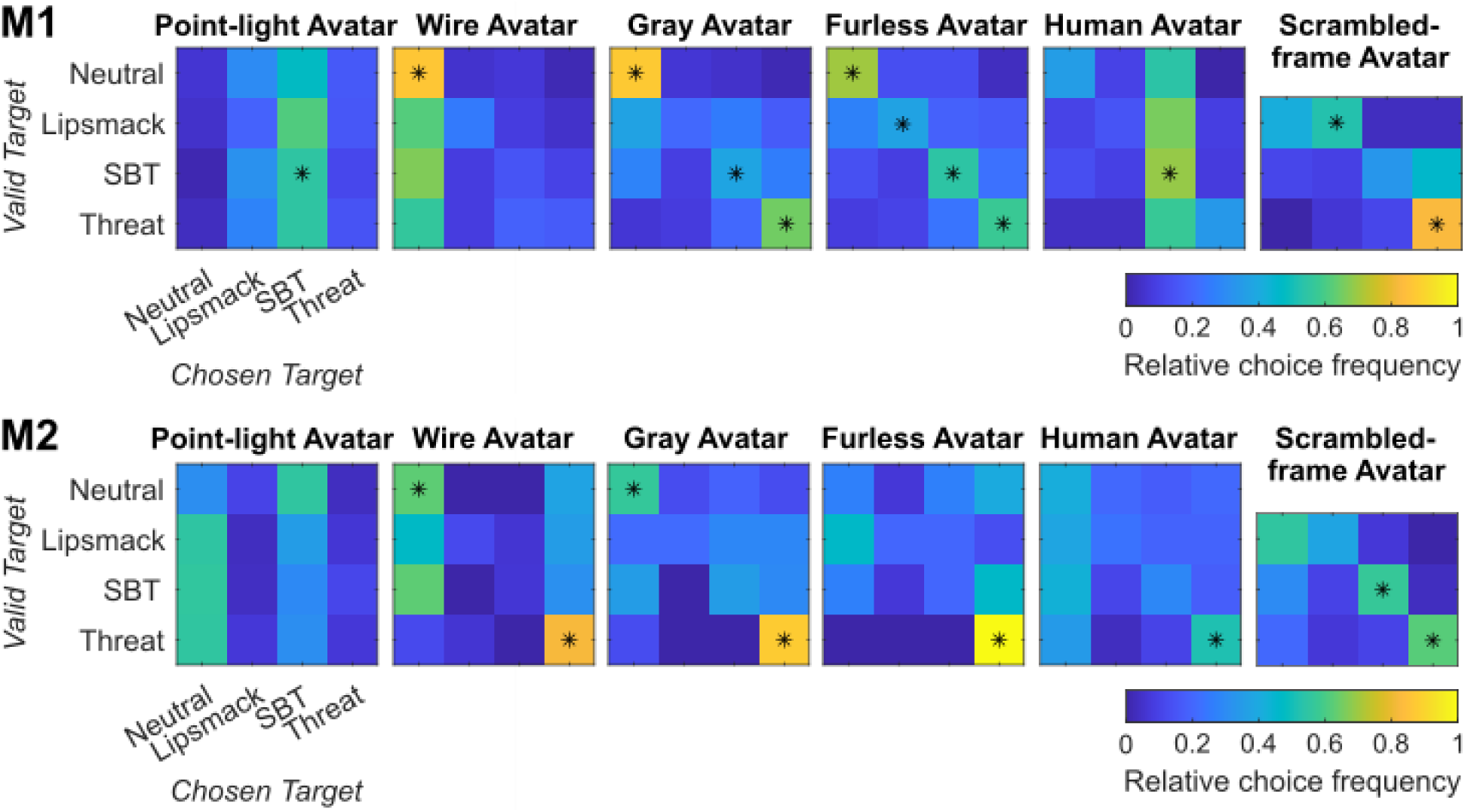
Expression categorization frequencies for avatar variants for monkey 1 (M1)/monkey 2 (M2). The first four avatar types increase in realism: point-light expressions (*N* per video = 60/29), wireframe mesh (*N* per video = 53/15), grayscale avatar (N=53/15) and furless avatar (*N* per video = 53/15). Additionally, a human avatar with monkey expressions (*N* per video = 59/29), and a naturalistic monkey avatar with time-scrambled frame order (*N* per video = 60/30) were tested (binomial tests, *p* < 0.05, FDR-corrected)

**Fig. S12.**
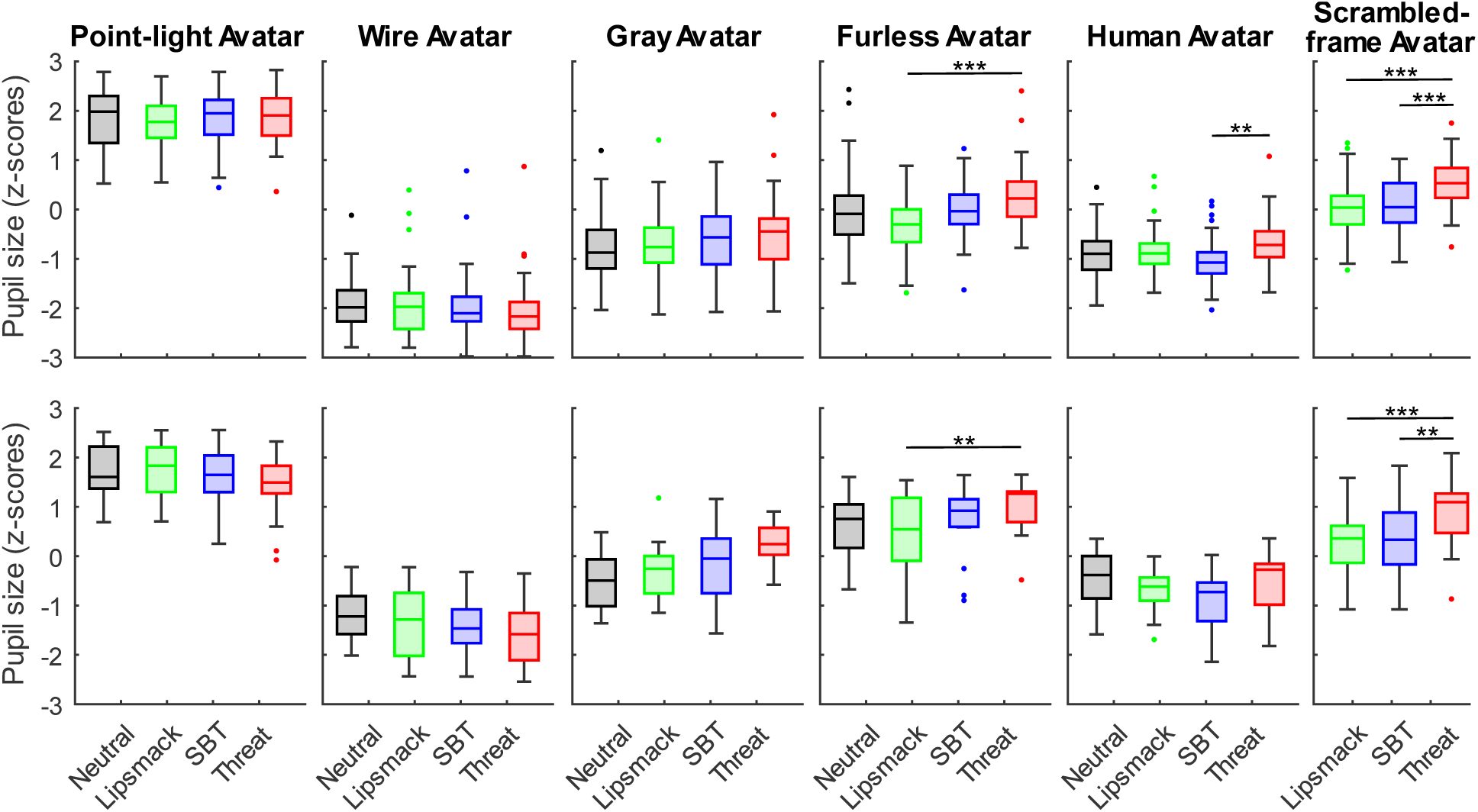
Boxplots of pupil sizes during the viewing of each expression category of the avatar variants for monkey 1 (M1) and monkey 2 (M2); \*\**p* < 0.01, \*\*\**p* < 0.001 (ANOVA, *post-hoc* Tukey-Kramer tests). The difference in pupil size of monkey 1 between the Neutral and the Threat category for the Gray Avatar marginally missed significance at *p* = 0.051. Systematic pupil size differences between avatar types are influenced by differences in stimulus type brightness

## References

Adams, R. B., & Kleck, R. E. (2003). Perceived Gaze Direction and the Processing of Facial Displays of Emotion. Psychological Science, 14(6), 644–647. doi:10.1046/j.0956-7976.2003.psci_1479.x

Adams, R. B., & Kleck, R. E. (2005). Effects of Direct and Averted Gaze on the Perception of Facially Communicated Emotion. Emotion, 5(1), 3–11. doi:10.1037/1528-3542.5.1.3

Andrew, R. J. (1963). Evolution of Facial Expression. Science, 142(3595), 1034–1041. doi:10.1126/science.142.3595.1034

Barrett, L. F., Adolphs, R., Marsella, S., Martinez, A. M., & Pollak, S. D. (2019). Emotional Expressions Reconsidered: Challenges to Inferring Emotion From Human Facial Movements. Psychological Science in the Public Interest, 20(1), 1–68. doi:10.1177/1529100619832930

Becker, D. V., Anderson, U. S., Mortensen, C. R., Neufeld, S. L., & Neel, R. (2011). The face in the crowd effect unconfounded: Happy faces, not angry faces, are more efficiently detected in single- and multiple-target visual search tasks. Journal of Experimental Psychology: General, 140(4), 637–659. doi:10.1037/a0024060

Beisner, B. A., & McCowan, B. (2014). Signaling context modulates social function of silent bared-teeth displays in rhesus macaques (Macaca mulatta). American Journal of Primatology, 76(2), 111–121. doi:10.1002/ajp.22214

Bergen, S., Huso, M. M., Duerr, A. E., Braham, M. A., Schmuecker, S., Miller, T. A., & Katzner, T. E. (2023). A review of supervised learning methods for classifying animal behavioural states from environmental features. Methods in Ecology and Evolution, 14(1), 189–202. doi:10.1111/2041-210X.14019

Bernstein, M., Erez, Y., Blank, I., & Yovel, G. (2018). An Integrated Neural Framework for Dynamic and Static Face Processing. Scientific Reports, 8(1), 7036. doi:10.1038/s41598-018-25405-9

Bernstein, M., & Yovel, G. (2015). Two neural pathways of face processing: A critical evaluation of current models. Neuroscience & Biobehavioral Reviews, 55, 536–546. doi:10.1016/j.neubiorev.2015.06.010

Blumrosen, G., Hawellek, D., & Pesaran, B. (2017). Towards Automated Recognition of Facial Expressions in Animal Models. Paper presented at the Proceedings of the IEEE International Conference on Computer Vision Workshops.

Breiman, L. (2001). Random Forests. Machine Learning, 45(1), 5–32. doi:10.1023/A:1010933404324

Clark, P. R., Waller, B. M., Burrows, A. M., Julle-Daniere, E., Agil, M., Engelhardt, A., & Micheletta, J. (2020). Morphological variants of silent bared-teeth displays have different social interaction outcomes in crested macaques (Macaca nigra). American Journal of Physical Anthropology, 173(3), 411–422. doi:10.1002/ajpa.24129

Crivelli, C., & Fridlund, A. J. (2019). Inside-Out: From Basic Emotions Theory to the Behavioral Ecology View. Journal of Nonverbal Behavior, 43(2), 161–194. doi:10.1007/s10919-019-00294-2

Cui, D., Sypré, L., Vissers, M., Sharma, S., Vogels, R., & Nelissen, K. (2023). Categorization learning induced changes in action representations in the macaque STS. Neuroimage, 265, 119780. doi:10.1016/j.neuroimage.2022.119780

Dal Monte, O., Costa, V. D., Noble, P. L., Murray, E. A., & Averbeck, B. B. (2015). Amygdala lesions in rhesus macaques decrease attention to threat. Nature Communications, 6(1), 10161. doi:10.1038/ncomms10161

Darwin, C. (1872). Expression of the emotions in man and animals (1 ed.). London, UK: John Murray.

de Waal, F. B. M., & Aureli, F. (1997). Conflict resolution and distress alleviation in monkeys and apes. In S. C. Carter, I. I. Lederhendler, & B. Kirkpatrick (Eds.), The Integrative Neurobiology of Affiliation (pp. 317–328). New York: New York Academy of Sciences.

de Waal, F. B. M., & Luttrell, L. M. (1985). The Formal Hierarchy of Rhesus Macaques - an Investigation of the Bared-Teeth Display. American Journal of Primatology, 9(2), 73–85. doi:10.1002/ajp.1350090202

Duchaine, B., & Yovel, G. (2015). A Revised Neural Framework for Face Processing. Annual Review of Vision Science, 1(Volume 1, 2015), 393–416. doi:10.1146/annurev-vision-082114-035518

Durán, J. I., & Fernández-Dols, J.-M. (2021). Do emotions result in their predicted facial expressions? A meta-analysis of studies on the co-occurrence of expression and emotion. Emotion, 21(7), 1550–1569. doi:10.1037/emo0001015

Ebitz, R. B., Pearson, J. M., & Platt, M. L. (2014). Pupil size and social vigilance in rhesus macaques. Frontiers in Neuroscience, 8, 100. doi:10.3389/fnins.2014.00100

Ekman, P. (1973). Universal facial expressions in emotion. Studia Psychologica, 15(2), 140.

Ekman, P. (1979). About brows: Emotional and conversational signals. In M. Von Cranach, K. Foppa, W. Lepenies, & D. Ploog (Eds.), Human Ethology: Claims and Limits of a New Discipline (pp. 169–202). Cambridge: Cambridge University Press.

Ekman, P. (1992). An argument for basic emotions. Cognition and Emotion, 6(3/4), 169–200. doi:10.1080/02699939208411068

Ekman, P. (2004). Emotional and conversational nonverbal signals. Paper presented at the Language, knowledge, and representation: Proceedings of the sixth international colloquium on cognitive science (ICCS-99).

Ekman, P., & Friesen, W. V. (1971). Constants across cultures in the face and emotion. Journal of Personality and Social Psychology, 17(2), 124–129. doi:10.1037/h0030377

Ekman, P., & Friesen, W. V. (1978). Facial action coding system: A technique for the measurement of facial movement. Palo Alto, CA: Consulting Psychologists Press.

Ekman, P., Friesen, W. V., & Hager, J. C. (2002). Facial action coding system - investigator’s guide. Salt Lake City, UT: Research Nexus.

Flack, J. C., & de Waal, F. (2007). Context modulates signal meaning in primate communication. Proceedings of the National Academy of Sciences, 104(5), 1581–1586. doi:10.1073/pnas.0603565104

Fortman, J. D., Hewett, T. A., & Halliday, L. C. (2001). The Laboratory Nunhuman Primate (1 ed.). Boca Raton: CRC Press.

Fridlund, A. J. (1994). Human facial expression: An evolutionary view. San Diego, CA: Academic Press.

Frijda, N. H., & Tcherkassof, A. (1997). Facial expressions as modes of action readiness. In J. A. Russell & J. M. Fernández-Dols (Eds.), The Psychology of Facial Expression (pp. 78–102). Cambridge: Cambridge University Press.

Frischen, A., Eastwood, J. D., & Smilek, D. (2008). Visual search for faces with emotional expressions. Psychological Bulletin, 134(5), 662–676. doi:10.1037/0033-2909.134.5.662

Giese, M. A., & Poggio, T. (2003). Neural mechanisms for the recognition of biological movements. Nature Reviews Neuroscience, 4(3), 179–192. doi:10.1038/nrn1057

Gong, M., & Smart, L. J. (2021). The anger superiority effect revisited: a visual crowding task. Cognition and Emotion, 35(2), 214–224. doi:10.1080/02699931.2020.1818552

Gothard, K. M., Battaglia, F. P., Erickson, C. A., Spitler, K. M., & Amaral, D. G. (2007). Neural Responses to Facial Expression and Face Identity in the Monkey Amygdala. Journal of Neurophysiology, 97(2), 1671–1683. doi:10.1152/jn.00714.2006

Gothard, K. M., Erickson, C. A., & Amaral, D. G. (2004). How do rhesus monkeys (Macaca mulatta) scan faces in a visual paired comparison task? Animal Cognition, 7, 25–36. doi:10.1007/s10071-003-0179-6

Grujic, N., Polania, R., & Burdakov, D. (2024). Neurobehavioral meaning of pupil size. Neuron, 112(20), 3381–3395. doi:10.1016/j.neuron.2024.05.029

Hadj-Bouziane, F., Bell, A. H., Knusten, T. A., Ungerleider, L. G., & Tootell, R. B. H. (2008). Perception of emotional expressions is independent of face selectivity in monkey inferior temporal cortex. Proceedings of the National Academy of Sciences, 105(14), 5591–5596. doi:10.1073/pnas.0800489105

Hasselmo, M. E., Rolls, E. T., & Baylis, G. C. (1989). The role of expression and identity in the face-selective responses of neurons in the temporal visual cortex of the monkey. Behavioural Brain Research, 32(3), 203–218. doi:10.1016/S0166-4328(89)80054-3

Hinde, R. A. (1966). Ritualization and social communication in Rhesus monkeys. Philosophical Transactions of the Royal Society of London. Series B, Biological Sciences, 251(772), 285–294. doi:doi:10.1098/rstb.1966.0012

Hoffman, K. L., Gothard, K. M., Schmid, Michael C., & Logothetis, N. K. (2007). Facial-Expression and Gaze-Selective Responses in the Monkey Amygdala. Current Biology, 17(9), 766–772. doi:10.1016/j.cub.2007.03.040

Izard, C. E. (1994). Innate and universal facial expressions: Evidence from developmental and cross-cultural research. Psychological Bulletin, 115(2), 288–299. doi:10.1037/0033-2909.115.2.288

Kanazawa, S. (1996). Recognition of facial expressions in a Japanese monkey (Macaca fuscata) and humans (Homo sapiens). Primates, 37(1), 25–38. doi:Doi 10.1007/Bf02382917

Kawai, N., Kubo, K., Masataka, N., & Hayakawa, S. (2016). Conserved evolutionary history for quick detection of threatening faces. Animal Cognition, 19(3), 655–660. doi:10.1007/s10071-015-0949-y

Landman, R., Sharma, J., Sur, M., & Desimone, R. (2014). Effect of distracting faces on visual selective attention in the monkey. Proceedings of the National Academy of Sciences, 111(50), 18037–18042. doi:doi:10.1073/pnas.1420167111

Le Mau, T., Hoemann, K., Lyons, S. H., Fugate, J. M. B., Brown, E. N., Gendron, M., & Barrett, L. F. (2021). Professional actors demonstrate variability, not stereotypical expressions, when portraying emotional states in photographs. Nature Communications, 12(1), 5037. doi:10.1038/s41467-021-25352-6

Lindquist, K. A., Gendron, M., Barrett, L. F., & Dickerson, B. C. (2014). Emotion perception, but not affect perception, is impaired with semantic memory loss. Emotion, 14(2), 375–387. doi:10.1037/a0035293

Maestripieri, D. (1997). Gestural Communication in Macaques: Usage and Meaning of Nonvocal Signals. Evolution of Communication, 1(2), 193–222. doi:10.1075/eoc.1.2.03mae

Mathis, A., Mamidanna, P., Cury, K. M., Abe, T., Murthy, V. N., Mathis, M. W., & Bethge, M. (2018). DeepLabCut: markerless pose estimation of user-defined body parts with deep learning. Nature Neuroscience, 21(9), 1281–1289. doi:10.1038/s41593-018-0209-y

Micheletta, J., Whitehouse, J., Parr, L. A., & Waller, B. M. (2015). Facial expression recognition in crested macaques (Macaca nigra). Animal Cognition, 18(4), 985–990. doi:10.1007/s10071-015-0867-z

Mielke, A., Waller, B. M., Perez, C., Rincon, A. V., Duboscq, J., & Micheletta, J. (2022). NetFACS: Using network science to understand facial communication systems. Behavior Research Methods, 54(4), 1912–1927. doi:10.3758/s13428-021-01692-5

Mori, M. (1970/2012). The uncanny valley (K. F. MacDorman & N. Kageki, Trans.). IEEE Robotics & Automation, 19(2), 3. doi:10.1109/MRA.2012.2192811

Morozov, A., Parr, L. A., Gothard, K., Paz, R., & Pryluk, R. (2021). Automatic Recognition of Macaque Facial Expressions for Detection of Affective States. eNeuro, 8(6). doi:10.1523/ENEURO.0117-21.2021

Mosher, C. P., Zimmerman, P. E., & Gothard, K. M. (2011). Videos of conspecifics elicit interactive looking patterns and facial expressions in monkeys. Behavioral Neuroscience, 125(4), 639–652. doi:10.1037/a0024264

Nahm, F. K. D., Perret, A., Amaral, D. G., & Albright, T. D. (1997). How do monkeys look at faces? Journal of Cognitive Neuroscience, 9(5), 611–623. doi:10.1162/jocn.1997.9.5.611

Nath, T., Mathis, A., Chen, A. C., Patel, A., Bethge, M., & Mathis, M. W. (2019). Using DeepLabCut for 3D markerless pose estimation across species and behaviors. Nature Protocols, 14(7), 2152–2176. doi:10.1038/s41596-019-0176-0

Nummenmaa, L., & Calvo, M. G. (2015). Dissociation between recognition and detection advantage for facial expressions: A meta-analysis. Emotion, 15(2), 243–256. doi:10.1037/emo0000042

Öhman, A., Lundqvist, D., & Esteves, F. (2001). The face in the crowd revisited: a threat advantage with schematic stimuli. Journal of Personality and Social Psychology, 80(3), 381–396. doi:10.1037/0022-3514.80.3.381

Parr, L. A., & Heintz, M. (2009). Facial expression recognition in rhesus monkeys, Macaca mulatta. Animal Behavior, 77(6), 1507–1513. doi:10.1016/j.anbehav.2009.02.024

Parr, L. A., Waller, B. M., Burrows, A. M., Gothard, K. M., & Vick, S. J. (2010). MaqFACS: A muscle-based facial movement coding system for the rhesus macaque. American Journal of Physical Anthropology, 143(4), 625–630. doi:10.1002/ajpa.21401

Partan, S. R. (2002). Single and Multichannel Signal Composition: Facial Expressions and Vocalizations of Rhesus Macaques (Macaca mulatta). Behaviour, 139(8), 993–1027. Retrieved from http://www.jstor.org/stable/4535968

Preuschoft, S. (1992). “Laughter” and “Smile” in Barbary Macaques (Macaca sylvanus). Ethology, 91(3), 220–236. doi:10.1111/j.1439-0310.1992.tb00864.x

Preuschoft, S. (2000). Primate Faces and Facial Expressions. Social Research, 67(1), 245–271. Retrieved from http://www.jstor.org/stable/40971384

Preuschoft, S., & van Hooff, J. A. (1995). Homologizing primate facial displays: a critical review of methods. Folia Primatologica, 65(3), 121–137. doi:10.1159/000156878

Pu, X., Fan, K., Chen, X., Ji, L., & Zhou, Z. (2015). Facial expression recognition from image sequences using twofold random forest classifier. Neurocomputing, 168, 1173–1180. doi:10.1016/j.neucom.2015.05.005

Rincon, A. V., Waller, B. M., Duboscq, J., Mielke, A., Pérez, C., Clark, P. R., & Micheletta, J. (2023). Higher social tolerance is associated with more complex facial behavior in macaques. Elife, 12(RP87008). doi:10.7554/eLife.87008

Russell, J. A. (1994). Is there universal recognition of emotion from facial expression? A review of the cross-cultural studies. Psychological Bulletin, 115(1), 102–141. doi:10.1037/0033-2909.115.1.102

Rychlowska, M., Miyamoto, Y., Matsumoto, D., Hess, U., Gilboa-Schechtman, E., Kamble, S., . . . Niedenthal, P. M. (2015). Heterogeneity of long-history migration explains cultural differences in reports of emotional expressivity and the functions of smiles. Proceedings of the National Academy of Sciences, 112(19), E2429–E2436. doi:doi:10.1073/pnas.1413661112

Sackett, G. P. (1966). Monkeys Reared in Isolation with Pictures as Visual Input: Evidence for an Innate Releasing Mechanism. Science, 154(3755), 1468–1473. doi:doi:10.1126/science.154.3755.1468

Savage, R. A., Lipp, O. V., Craig, B. M., Becker, S. I., & Horstmann, G. (2013). In search of the emotional face: Anger versus happiness superiority in visual search. Emotion, 13(4), 758–768. doi:10.1037/a0031970

Siebert, R., Taubert, N., Spadacenta, S., Dicke, P. W., Giese, M. A., & Thier, P. (2020). A Naturalistic Dynamic Monkey Head Avatar Elicits Species-Typical Reactions and Overcomes the Uncanny Valley. eNeuro, 7(4). doi:10.1523/ENEURO.0524-19.2020

Stettler, M., Lappe, A., & Giese, M. A. (2026). Facial expression recognition based on multi-domain norm-referenced encoding. Neural Networks, 197, 108384. doi:10.1016/j.neunet.2025.108384

Susskind, J. M., Lee, D. H., Cusi, A., Feiman, R., Grabski, W., & Anderson, A. K. (2008). Expressing fear enhances sensory acquisition. Nature Neuroscience, 11(7), 843–850. doi:10.1038/nn.2138

Taubert, J., & Japee, S. (2024). Real Face Value: The Processing of Naturalistic Facial Expressions in the Macaque Inferior Temporal Cortex. Journal of Cognitive Neuroscience, 36(12), 2725–2741. doi:10.1162/jocn_a_02108

Taubert, J., Japee, S., Murphy, A. P., Tardiff, C. T., Koele, E. A., Kumar, S., . . . Ungerleider, L. G. (2020). Parallel Processing of Facial Expression and Head Orientation in the Macaque Brain. Journal of Neuroscience, 40(42), 8119–8131. doi:10.1523/jneurosci.0524-20.2020

Taubert, N., Stettler, M., Siebert, R., Spadacenta, S., Sting, L., Dicke, P., . . . Giese, M. A. (2021). Shape-invariant encoding of dynamic primate facial expressions in human perception. Elife, 10. doi:10.7554/eLife.61197

Thierry, B., Demaria, C., Preuschoft, S., & Desportes, C. (1989). Structural convergence between silent bared-teeth display and relaxed open-mouth display in the Tonkean macaque (Macaca tonkeana). Folia Primatologica, 52(3-4), 178–184. doi:10.1159/000156396

Valente, D., Theurel, A., & Gentaz, E. (2018). The role of visual experience in the production of emotional facial expressions by blind people: a review. Psychonomic Bulletin & Review, 25(2), 483–497. doi:10.3758/s13423-017-1338-0

van Hooff, J. A. R. A. M. (1967). The Facial Displays of the Catarrhine Monkeys and Apes. In D. Morris (Ed.), Primate Ethology (pp. 7–68). Chicago, IL: Aldine Pub.

Vangeneugden, J., Vancleef, K., Jaeggli, T., VanGool, L., & Vogels, R. (2010). Discrimination of locomotion direction in impoverished displays of walkers by macaque monkeys. Journal of Vision, 10(4), 22, 21–19. doi:10.1167/10.4.22

Vlaeyen, J. M. R., Heesen, R., Kret, M. E., Clay, Z., Bionda, T., & Kim, Y. (2022). Bared-teeth displays in bonobos (Pan paniscus): An assessment of the power asymmetry hypothesis. American Journal of Primatology, 84(9), e23419. doi:10.1002/ajp.23419

Waller, B. M., Whitehouse, J., & Micheletta, J. (2016). Macaques can predict social outcomes from facial expressions. Animal Cognition, 19(5), 1031–1036. doi:10.1007/s10071-016-0992-3

Waller, B. M., Whitehouse, J., & Micheletta, J. (2017). Rethinking primate facial expression: A predictive framework. Neuroscience & Biobehavioral Reviews, 82, 13–21. doi:10.1016/j.neubiorev.2016.09.005

Wehrheim, M., Alamooti, S. T., Ramezanpour, H., & Kar, K. (2025). Facial expression discrimination emerges from neural subspaces shared with detection and identity. bioRxiv, 2025.2008.2025.672186. doi:10.1101/2025.08.25.672186

Wolfensohn, S., & Lloyd, M. (2003). Handbook of Laboratory Animal Management and Welfare (3 ed.). Oxford, UK: Blackwell Publishing Ltd.

Yang, Z., & Freiwald, W. A. (2021). Joint encoding of facial identity, orientation, gaze, and expression in the middle dorsal face area. Proceedings of the National Academy of Sciences, 118(33), e2108283118. doi:10.1073/pnas.2108283118

Zhang, J., Sato, W., Kawamura, N., Shimokawa, K., Tang, B., & Nakamura, Y. (2024). Sensing emotional valence and arousal dynamics through automated facial action unit analysis. Scientific Reports, 14(1), 19563. doi:10.1038/s41598-024-70563-8

Zhu, Q., Nelissen, K., Van den Stock, J., De Winter, F.-L., Pauwels, K., de Gelder, B., . . . Vandenbulcke, M. (2013). Dissimilar processing of emotional facial expressions in human and monkey temporal cortex. Neuroimage, 66, 402–411. doi:10.1016/j.neuroimage.2012.10.083

